# Neuroligin 3 highlights sexually dimorphic circuitry in Drosophila social spacing

**DOI:** 10.1101/2024.10.28.620699

**Authors:** JW Robinson, AT Bechard, MR Evans, R Ataei, J Kurbaj, K Mosuro, JR Isaacson, S Pillay, DS Lin, A Sahota, JN de Belle, GI Robinson, AJ Moehring, AF Simon

## Abstract

In *Drosophila melanogaster*, the autism-related Neuroligin 3 (Nlg3) protein is a postsynaptic membrane protein important for synapse development and regulation, which plays a role in social spacing behaviour. Here, we report the localization of Nlg3 to the calyx of the mushroom bodies (MB), optic lobes (OL), and protocerebral bridge (PB). Using RNA interference, *nlg3* knockdown in each of these structures recapitulated the effect of knocking down it in all *nlg-3* neurons. Hyperactivation and silencing of these neurons in the MB, but not the PB, controls social space in males and females, while hyperactivating and silencing of all *nlg3*-expressing neurons, including within the MB, PB, and OL, regulates male and female social space. Knocking down neurotransmitter biosynthesis enzymes, which decreases the amount of neurotransmitter release, showed that reducing acetylcholine release from the MB decreased female social space, whereas knocking down any dopamine receptor in the MB increased male social space. Lastly, to investigate the sexually dimorphic effects on social spacing previously seen in *nlg3* mutants, we examined a subset of sexually dimorphic *fruitless-*expressing (*fru)P1* neurons known to regulate sexually dimorphic behaviours. Hyperactivation of those *fruP1* neurons decreased social space in both sexes, while silencing those *fruP1* neurons specifically increased male social space without affecting females. Our findings highlight a sex-specific social space neural circuitry that includes the OL, MB, and *fruP1* neurons, while uncovering the underlying basis of some of the sex differences in this behaviour.

**Article Summary:** In vinegar flies (*Drosophila melanogaster*), the autism-related Neuroligin 3 protein (Nlg3) controls neuronal development and regulation but also affects fly social behaviour. Nlg3 is localized to the mushroom bodies (MB), protocerebral bridge, and the optic lobes. We show that those structures are important in determining social space in a sex dependant manner. In addition, reducing acetylcholine release from the MB affects female social space, while reducing dopamine receptors of the MB only affect male behaviour. Finally, the *fruitless* sexually dimorphic neurons control social behavior differently in males and females.

## Introduction

Social behaviours are fundamental to the interaction and communication within a community and are influenced by environmental cues which are processed and integrated in the brain. Abnormal social behaviours, such as difficulties with communication or interactions with others, are clinical characteristics of individuals with autism spectrum disorders (ASD). Using *Drosophila melanogaster,* we can dissect the neural and genetic underpinnings of social interactions in a simplified system (Sokolowski 2001).

Flies exhibit social behaviours that, while simpler, can reflect the basic principles of social interaction found in more complex organisms (Sokolowski 2010). By examining how environmental cues are integrated and influence social behaviours in Drosophila, we can gain insights into the mechanisms that might be disrupted in ASD, providing a basis for understanding the molecular and neural circuitries involved in these disorders (Ferreira and Moita 2019).

An important component of Drosophila social interactions within a group context is social spacing – the distance of one individual to another in a group. We use the social space assay to assess the outcome of group behaviour displayed by flies as a measure of the average number of flies within four-body lengths (4BL) in the social space chamber (Simon et al. 2012). We and other research groups have used this assay as a metric for behavioural analysis to understand and reveal the basic underlying neural circuitry involved (Burg et al. 2013; Fernandez et al. 2017; Xie et al. 2018; Yost et al. 2020).

So far, we know that mutations in genes affecting eye function, have been found to make flies less social (Simon et al. 2012; Burg et al. 2013; Jiang et al. 2020). Both males and females have decreased social space with mutants associated to fly gustation and mechanosensation (Jiang et al. 2020). In contrast, *Orco* mutants, which affect olfactory perception, do not appear to influence social spacing in males, but result in decreased social space in females. Together, these suggest a sensory modality-specific genetic influence on social behaviour that can have sex-specific effects (Simon et al. 2012; Jiang et al. 2020).

Other genes expressed in the brain have also been shown to influence Drosophila social behaviour, such as genes important for synaptic transmission and synaptogenesis (Burg et al. 2013; Wise et al. 2015; Fernandez et al. 2017; Ueoka et al. 2018; Brenman-Suttner et al. 2019; Hope et al. 2019; Jiang et al. 2020; Kanellopoulos et al. 2020; Yost et al. 2020; Shilpa et al. 2021; Cao et al. 2022). At the synapse level, the concentration of neurotransmitters, which is tightly regulated by biosynthesis enzymes or vesicular packaging proteins, have been shown to affect social spacing. Tyrosine hydroxylase (TH), the rate-limiting enzyme in dopamine synthesis, and the vesicular monoamine transporter (VMAT), which packages dopamine into synaptic vesicles, both regulate dopamine levels (Simon et al. 2009; Lawal et al. 2010). Modulating the abundance or activity of these proteins influences social spacing in Drosophila, indicating that dopamine plays a role in regulating social space (Fernandez et al. 2017).

Many human homologs of neural genes have also emerged as potential candidates in neuropsychiatric disorders, bridging the gap between basic biological processes and complex human pathologies (Grant 2012). The gene *neurobeachin*, which is implicated in ASD in humans, encodes a structural scaffolding postsynaptic density protein (Castermans et al. 2003). In Drosophila, mutations in the homolog *rugose* result in altered fly social spacing, indicating a conservation of function across species (Wise et al. 2015). Furthermore, synaptic proteins Neuroligin and Neurexin, which facilitate neurotransmission by aiding synapse formation and receptor recruitment, have been shown to influence social behaviour (Hahn et al. 2013). In humans, *Neuroligin* gene homologs are correlated with ASD (Jamain et al. 2003; Südhof 2008). Similarly, in Drosophila, mutations in these genes lead to altered social behaviours, further revealing the genetic underpinnings of social space (Corthals et al. 2017).

Drosophila Neuroligin (Nlg) is a family of post-synaptic membrane proteins with the conserved function of synapse formation and maintenance by interacting with presynaptic Neurexin to mediate cell-cell adhesion and facilitate neurotransmission (Südhof 2008). During Drosophila neural development, larval *nlg3* mutants have a higher number of synaptic connections but less GluRIIA recruitment and synaptic transmission compared to control flies (Xing et al. 2014). Furthermore, *nlg3* knockouts display a sexually dimorphic and age-associated effect on fly aggression and social space (Yost et al. 2020). Mutant *nlg3* flies had lower climbing performance in both males and females by over 30%, however, *nlg3* deletion decreases only male, but not female aggression; while increasing male social space in 3-4 day old flies, and in both sexes by 7-10 days old (Yost et al. 2020).

The mushroom bodies (MB), also known to play a role in Drosophila social spacing, are a well-studied invertebrate brain structure primarily involved in fly olfactory learning and memory. The MB are comprised of approximately 2,500 neurons and its unusually structured shape suggests a highly specialized organization and function (Erber et al. 1987). The MB have a flexible organization in response to social isolation and are the central processing structure of the Drosophila brain (Technau 1984). Consisting of densely packed neurons called Kenyon cells, the MB integrates many sources of sensory information to modulate fly behavioural responses. Social space assays have been conducted on flies with a mutation in an ion channel selectively targeted in the MB and lead to decreased social space, observing that the cholinergic signaling on its own can rescue this altered phenotype in social spacing (Burg et al. 2013).

Sexually dimorphic behaviours in Drosophila such as male courtship, female receptivity, and social interactions are known to be regulated by *fruitless* (*fru*) neurons (Pan and Baker 2014; Brenman-Suttner et al. 2018; Chowdhury et al. 2020). The *fruitless* gene encodes a transcription factor important for sex determination in Drosophila development and produces male-specific transcripts that are essential for the development and function of male-specific neural circuits (Sato and Yamamoto 2020). Regulated by multiple promoters each expressing unique transcripts, the *fruP1* transcript undergoes sex-specific splicing and contributes to the sexual differentiation of neural circuits influencing behaviours such as courtship and aggression (Wiedemann 2007; Baker et al. 2024). Promoters *p2-p5* are necessary for adult viability and fly morphology that is however not sex specific (Ito et al. 1996). Neurons expressing *fru* are widespread interneurons in the adult fly brain and ventral nerve cord, where fast acting *fru* neurons release acetylcholine, GABA, and glutamate, while other *fru* neurons also release dopamine, serotonin, tyramine, and octopamine (Palmateer et al. 2023).

While progress has been made in identifying individual genes, synaptic proteins, and neuropils like the MB that influence Drosophila social spacing, the integration of molecular players within precise circuitry or other brain structures mediating social spacing behaviour are not fully understood. This research looks deeper into the role *nlg3* has on Drosophila social space. Based on Nlg3 localization and the sex-specific differences observed, we elucidated part of the neural circuitry that directly influences social spacing behaviour.

## Materials and Methods

### Fly Stocks and Husbandry

All flies were reared mixed sex in bottles containing our own fly food recipe (brown sugar, corn meal, yeast, agar, benzoic acid, methyl paraben, and propionic acid). Rearing conditions were controlled during development and aging at 50% humidity, 25⁰C, and on a 12:12 hour light:dark cycle. Parents were removed after seven days of egg laying resulting in a maximum parental age of 14 days, as parental age affects social spacing of the offspring (Brenman-Suttner et al. 2018). Canton-S (Cs) control line was initially sourced from Seymour Benzer in 1998. UAS-effector lines were outcrossed five times into the Cs control line to minimize behavioural variation due to genetic background. All genotypes used are reported in **Supp. Table 1.** A generalized crossing scheme for both the V10 and Cs backgrounds used to create the appropriate experimental treatment and genetic controls can be found in the supplementary material (**Supp. Fig. 1**). The results obtained including those UAS effector lines, despite being in a different genetic background, are presented in **Supp. Fig. 8-11** where the effect of genetic background may be observed in some experiments.

### Generation of a UAS-nlg3 RNAi line

The shRNA sequences targeting nlg3 were obtained by using the model provided by Vert et al. (2006). The length of the *nlg3* shRNA was 60 bases for the full hairpin; and was ∼7.8kbp for the entire pVALIUM20 plasmid containing the shRNA was ∼7.8kbp. The hairpin sequence used was:

### CAGCAACTCGAACTCGAACTAtagttatattcaagcataTAGTTCGAGTTCGAGTTGCTG

Within the hairpin sequence, uppercase letters represent the nucleotides that bind to the *nlg3* mRNA, while lowercase letters represent the nucleotides that connect the two halves of the pin. The shRNA constructs were cloned into the vector pVALIUM20 (vermillion eye color maker, (Ni et al. 2011) using the protocol in Chang et al. (2014). DNA was injected into genotype *y v, nanos-integrase, attP40/attP40* (Bloomington Stock #: 25709) embryos and were mated to the parental line after reaching adulthood. Then, offspring containing the transgene were backcrossed five times into the Cs control background (Vert et al. 2006; Chang et al. 2014).

### RT-qPCR to confirm effect of the UAS-nlg3 RNAi line

Heads from ∼20 male or female flies (3–5 days old) were collected separately. Control and RNAi treatment groups were selected to match the genotypes and ages used in the behavioural assays, ensuring consistency across experiments. Total RNA was extracted using TRIzol reagent (#15596026-Invitrogen) according to the manufacturer’s instructions. RNA samples were treated with DNase (TURBO^TM^ DNase-Invitrogen) to eliminate genomic DNA contamination, and cDNA was synthesized using the cDNA Synthesis Kit (#1708841-BIO-RAD). RT-qPCR was performed using the PowerUp^TM^ SYBR^TM^ Green Master Mix (Thermo Fisher Scientific) on a CFX Connect Real-Time PCR Detection System (BIO-RAD). Relative expression levels were calculated using the ΔΔCq method, with *Tubulin84* serving as the internal control gene. The qPCR analysis confirmed that nlg3 transcript levels were significantly reduced in RNAi lines compared to controls. Knockdown efficiency was observed in both sexes, with females showing stronger suppression than males, as shown in **Supp. Fig. 2**.

### Immunocytochemistry

#### Anti-Nlg3

Whole brain dissections on 3 to 4 day old, male or female brains were fixed within 20 minutes of dissection in Bouin’s solution. Following fixation and washing, brains were blocked in 5% NGS in 1X PBS and stained with guinea pig anti-Nlg3 primary antibody (1:1000, Invitrogen) overnight at 4 °C, which is a polyclonal primary antibody kindly provided by Dr. Brian Mozer (Yost et al. 2020). After washing, brains were stained with goat anti-guinea pig Alexa 488 (1:500, Invitrogen) for 2 hours at room temperature. Next, all samples were left in glycerol for 1 hour then were mounted on slides using Fluoroshield^TM^ media, stored at 4 °C and imaged the following day. Fluorescent images were visualized with a AxioImager Z1 Compound Fluorescent microscope (Zeiss). Note: we used the entirety of our polyclonal antibody stock and have been unsuccessful in generating another functioning antibody for Nlg3. We switched to genetic constructs that tag Nlg3 with a GFP reporter (*MiMIC-nlg3* and *nlg3-Gal4 > mCD8-GFP*) that both verify our findings.

#### Anti-GFP

Whole brain dissections on 3 to 4 day old males or females were fixed within 20 minutes of dissection in 4% paraformaldehyde in 1X PBS-T (0.3% Triton-X). Following fixation and washing, brains were blocked in 5% normal goat serum (NGS) in PBS-T and stained with rabbit anti-GFP (1:1000, Invitrogen) and mouse anti-Brp (nc82; 1:30, Developmental Studies Hybridoma Bank) for 48 hours at 4 °C. After washing, brains were stained with Alexa 660 anti-mouse (1:400, Invitrogen) and Alexa 488 anti-rabbit (1:800, Invitrogen) for 48 hours at 4°C. Samples were mounted on slides in Fluoroshield^TM^ media, stored at 4 °C and imaged the following day. Slides were imaged on an SP8 Lightening Confocal microscopy system (Leica). All images were visualized with LAS X 2.0 software (Leica).

### Social Space Assay

The assay was performed as previously described (McNeil et al. 2015; Yost et al. 2020). In short, flies were collected after eclosion and group housed until aged at 3 to 4 days old. Yost et al. (2024) have shown that the flies are no longer virgin after the age treatment we use for these experiments. Twenty-four hours prior to the assay, flies were sexed under cold anesthesia and separated into vials with food containing 12-17 flies for each treatment tested. Two hours before the assay, each treatment was transferred into new vials and acclimated to standard conditions (25°C and 50% humidity). Ten minutes prior to test start, if the standard acclimation temperature is different than testing condition temperature, treatments were set to the testing temperature required for neuronal hyperactivation (*TrpA1* at 27°C and 50% humidity) or silencing (*Shi^ts^* at 29°C and 50% humidity). For those experiments we compared the effect of the treatment at higher temperature (activation of genetic constructs and alteration of neural transmission) to the typical room testing temperatures (25°C, control: inactivated genetic constructs and basal transmission of neurons). So, the same genotypes were tested at 2 different temperatures. All genetic background lines used to create the experimental lines displayed no effect of the temperature changes (**Fig. 2**). At the time of testing, flies were loaded into the triangular chamber and knocked to the bottom of the chamber to induce a startle response that forces the flies into a group at the top of the chamber within the first minute. Flies are then given 20-40 minutes to settle at their preferred social space. Each treatment was loaded into the social space apparatus and photos were taken as soon as flies have settled as a group (McNeil et al. 2015). Social space assay photos were processed with ImageJ (National Institute of Health, Bethesda, MA, USA) to calculate the number of flies within a four-body length (4BL or ∼1cm) from macros (as described in Yost et al. 2020). Image analysis data were imported to GraphPad Prism 10 software for statistical analysis and graphical representation.

### Climbing Assay

The assay was performed as previously described (McNeil et al. 2015; Yost et al. 2020). In this assay we are testing flies’ stamina, ability to respond to startle, and locomotion. Any lines with flies able to reach the top vial (performance index, PI, more than zero) in 15 seconds do not have impairments preventing testing in the social space assays (McNeil et al. 2015; Yost et al. 2020). Flies were aged to 2 to 4 days old and 24 hours prior to conducting the assay were sexed under cold anesthesia and separated in vials with food containing 40 flies for each treatment. Two hours before the assay, each treatment was transferred into new vials and acclimated to testing conditions (25⁰C and 50% humidity). Ten minutes before the assay, if the standard acclimation temperature is different than testing condition temperature, treatments were set to testing temperature required for neuronal hyperactivation (*TrpA1* at 27°C and 50% humidity) or silencing (*Shi^ts^*at 29°C and 50% humidity). Each treatment was transferred into a testing tube and inserted into the climbing assay. Flies were forced to the bottom of the tube and given 15 seconds to climb to the tube on top of the apparatus. This timeframe was chosen to allow approximately 80% of control flies to reach the top vial. After 15 seconds, the apparatus was closed and the number of flies in the top and bottom tubes were counted to calculate a performance index of percent flies that reached the top vial.

### Statistical Analysis

Statistical analysis and graphical representation were performed in GraphPad Prism (RRID:SCR_002798, version 7.0a for Mac, GraphPad Software, La Jolla, CA, USA, https://www.graphpad.com). Normality test was used to verify normality and homoscedasticity test was used to verify the homoscedasticity of the distribution before performing an ANOVA test. Two-way ANOVAs were performed on data presented on the same graph, which were collected simultaneously and had 2 variables in 2 groups. Following the ANOVAs, Šídák’s post-hoc tests were performed within each sex to compare between different genotypes or transmission states, while correcting for multiple comparisons. The results of all the comparison tests for social space analysis can be found in **Supp. Table 2.** Supplemental figures have all ANOVA results and comparison tests in **Supp. Table 3.** P-values are reported on graphs if *p* < 0.05.

### Drawings

Drawings presented in figures were created on the app Procreate 5.3.5 (Savage Interactive Pty Ltd, Hobart, Australia).

## Results

### Neuroligin 3 protein is expressed at low levels in the calyx of the mushroom bodies, the protocerebral bridge, and the optic lobes

We investigated the expression patterns of the Nlg3 protein in the adult brain using an anti-Nlg3 polyclonal antibody. We found Nlg3 protein was localized within the calyx (Ca) of the mushroom bodies (MB), the protocerebral bridge (PB), and the optic lobes (OL) all in low levels compared to the background autofluorescence of brain lipids (**Fig. 1A,B**). For confirmation of the findings done with our antibody, we used two genetic constructs (*MiMIC* p-element insertion within *nlg3*, as well as an *nlg3-Gal4* insertion within the gene – see **Supp. Table 1** for specific localization). The *MiMIC-nlg3* insertion which incorporates a GFP reporter into the coding sequence of the host gene *nlg3* (Venken et al. 2011). From this construct we saw GFP expressed at low levels to the MB and OL and at very low fluorescence we can see the PB when compared to background autofluorescence of the Drosophila brain (**Fig. 1C,D**). Confirmation of expression in the calyx and PB was observed with the *nlg3-Gal4* insertion driving *mCD8-GFP* at higher levels than endogenous *nlg3* (**Fig. 1E,F**). Male and female localization of Nlg3 did not appear to be different either qualitatively or quantitatively in 3-4 day old adult flies (representative images of only the males are shown). Next, using the *nlg3-Gal4* driver to express GFP in **Fig. 1E** and **1F**, we activated an RNAi specific to *nlg3* (outcrossed in the Cs background) to all cells that express *nlg3*. In both males and females, both genetic control fly lines (*UAS/+* and *RNAi/+*) did not have different social space compared to the Cs genetic background control strain (**Fig. 1G,H**). However, in both males and females again, we did see a decrease in the number of flies within 4BL for the RNAi knocking in *nlg3*-expressing cells (One-Way ANOVA *p* < 0.0001, post-hoc comparison of ngl3>UAS-RNAi nlg3 to other controls *p* < 0.0001, see details in **Fig. 1G,H**).

**Figure 1.**
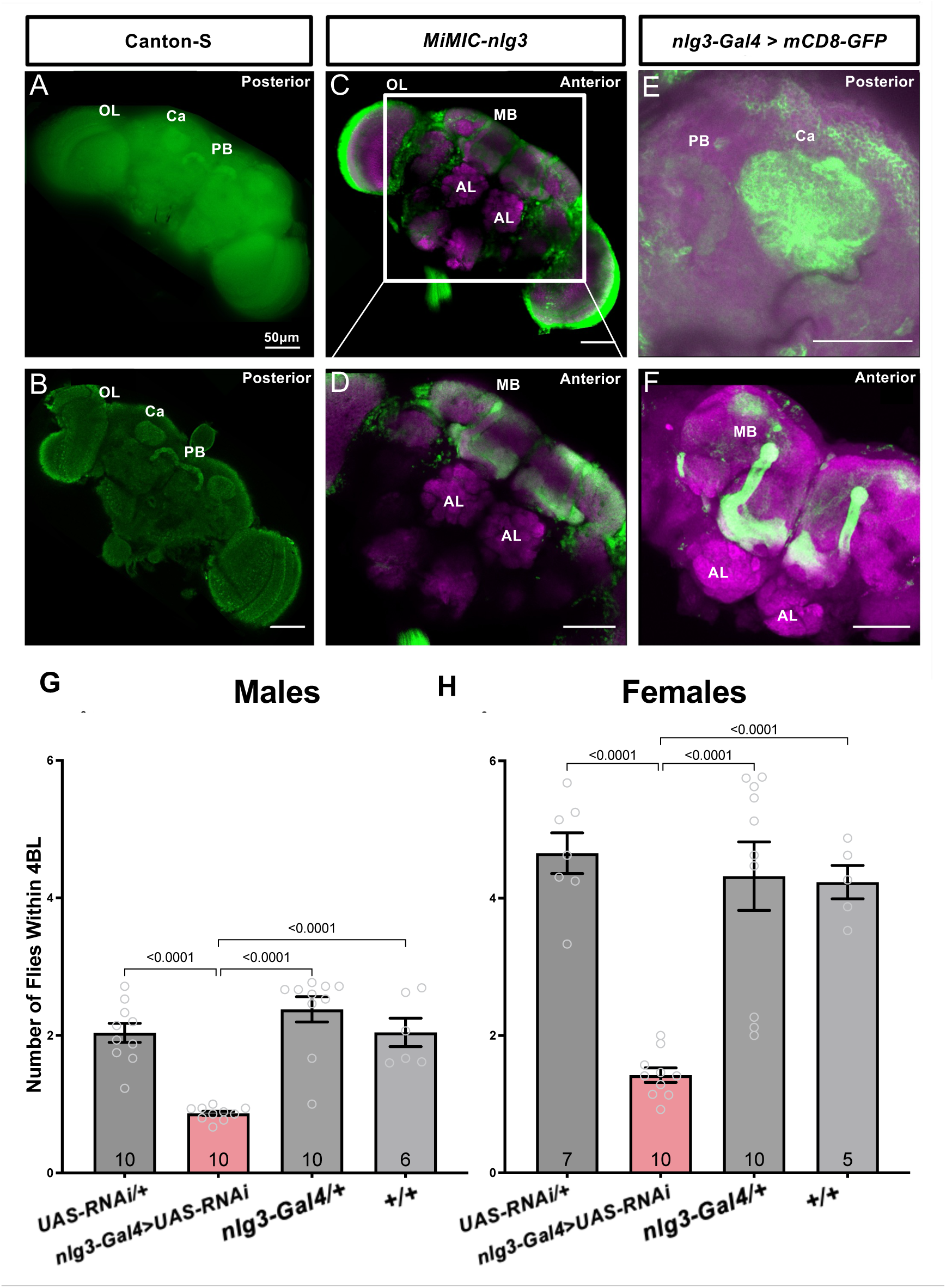
Nlg3 is expressed in the calyx of the mushroom bodies, throughout the protocerebral bridge, and in the optic lobes, and that expression is important for social space. **A-B:** Immunofluorescent imaging of polyclonal antibody specific to the c-terminal intracellular domain with anti-Nlg3 (green) on Canton-S male fly whole brains dissected at 3-4 days old. Magnification: 20X. **A.** Fluorescent imaging on posterior side of adult brain shows three main structures expressing *nlg3* in low levels: calyx of the MB (Ca), protocerebral bridge (PB), and the optic lobes (OL). **B.** Confocal imaging on posterior side of adult brain shows definition of Nlg3 expression in the Ca, PB, and OL (medulla and lamina). **C-D**: Expression patterns of MiMIC construct (*Mi{y[+mDint2]=MIC}nlg3[MI08924]*) incorporating *mCD8-GFP* into *nlg3*. Immunofluorescence with anti-GFP (green) and nc82 (magenta). **C.** Confocal imaging on anterior side of adult brain at 20X magnification indicates low expression of Nlg3 in the MB and OL. **D.** Confocal imaging at 40X magnification highlighting expression in the MB. E-F: Expression patters of *MiMIC nlg3-Gal4* genetic construct (*Mi{Trojan-Gal4}nlg3>UAS-mCD8::GFP*) driving exogenous expression of *mCD8-GF*P with *nlg3*. Immunofluorescence with anti-GFP (green) and nc82 (magenta). **E.** Confocal imaging of posterior side of adult brain at 40X magnification indicates low expression patterns in PB and Ca. **F.** Confocal imaging of anterior side of adult brain at 40X magnification shows expression throughout the MB. Antennal Lobes show for reference (AL). Scale bar, 50µm. **G-H**: Social space of flies with RNAi knockdown of *nlg3* targeted specifically to all *nlg3*-expressing cells. There is a decrease in the average number of flies within 4 body lengths (4BL) in male (G; one-way ANOVA*: F_3,32_* = 1.771, *p* = 0.0003, post-hoc Šidák’s test p<0.0001 when comparing *nlg3*>UAS-RNAi *nlg3* to all controls) and female (H; one-way ANOVA*: F_3,28_* = 3.967, *p* < 0.0001, post-hoc Šidák’s test p<0.0001 when comparing *nlg3*>UAS-RNAi *nlg3* to all controls).

### Neuroligin 3 expression in the calyx of the mushroom bodies, the protocerebral bridge, and the optic lobes is required for normal social spacing

To determine the behavioural function of Nlg3 located in these structures, we targeted RNAi knock down of *nlg3* independently in each of the MB, PB, and OL and tested fly social space. We observed a behavioural effect of knocking down *nlg3* in the MB (**Fig. 2A**; Two-way ANOVA – Effect of genotype: *F_1,39_* = 14.71, *p* = 0.0004), in the PB (**Fig. 2B**; Two-way ANOVA – Effect of genotype: *F_1,40_* = 19.02, *p* < 0.0001; Šídák’s multiple comparisons test p = 0.001), and the OL (**Fig. 2C**; Two-way ANOVA – Effect of genotype: *F_1,31_* = 7.191, *p* = 0.0116). Then, we used a climbing assay to assess fly locomotion and their ability to respond to a startle response and climb; all to ensure the treatment flies could perform in the social space assay. While there was no reduction in climbing when knocking down *nlg3* in the MB (**Supp. Fig. 3A**), there was a reduction in climbing when knocking down *nlg3* in each of the PB and OL (**Supp. Fig. 3B,C**). The flies were able to climb and therefore were able to perform the social space assay. We also observed the variations in climbing are not correlated to the variations in social spacing.

**Figure 2.**
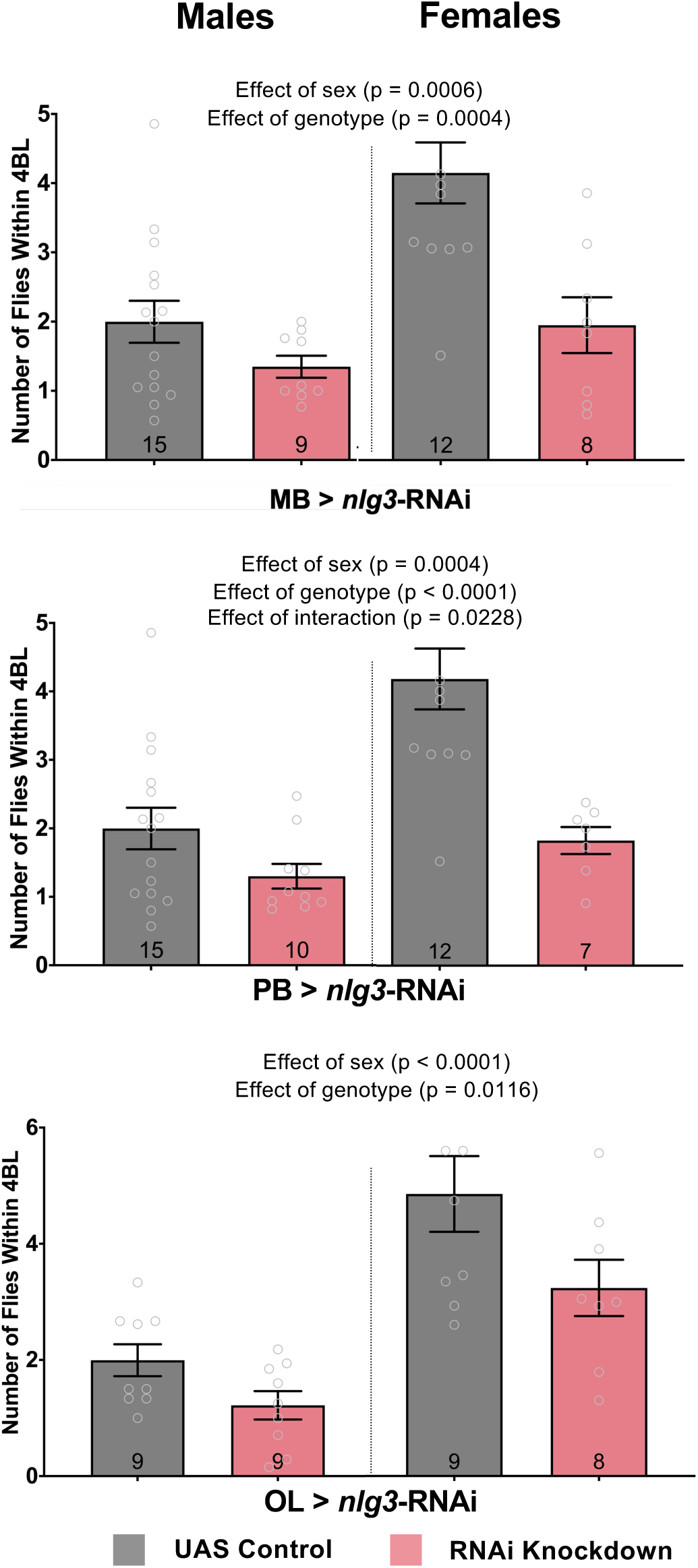
Knocking down Nlg3 specifically in each of the mushroom bodies, the protocerebral bridge or the optic lobes specifically increases fly social space in males and females. **A-C:** Average number of flies within 4 body lengths (4BL) in male and female flies with RNAi knockdown of *nlg3* targeted specifically to the mushroom bodies (A), protocerebral bridge (B), and the optic lobes (C). **A.** The number of flies within 4BL decreased in both males and females in flies with *nlg3* knockdown targeted to the MB (two-way ANOVA – Effect of genotype*: F_1,39_* = 14.71, *p* = 0.0004). **B.** The average number of flies within 4BL decreased in both males and females in flies with *nlg3* knockdown targeted to the PB (two-way ANOVA – Effect of genotype: *F_1,40_* = 19.02, *p* < 0.0001). **C.** Knocking down *nlg3* in the OL decreases the number of flies within 4BL in both males and females compared to genetic control flies (two-way ANOVA – Effect of genotype: *F_1,31_* = 7.191, *p* = 0.0116). Grey colour: genetic control *driver/+*; pink colour: *driver/RNAi*. The results of all two-way ANOVA and *post-hoc* tests performed can be found in **Supp. Table 2.** N = 7 – 14 for all treatments. Error bars are +/-s.e.m.

### Hyperactivation and silencing nlg3-expressing neurons, independently or simultaneously, changes fly social space in a sex-specific manner

Next, we looked at genetically manipulating neural transmission within the brain structures where Nlg3 is localized (**Fig. 1**) to examine their effect on social space. When hyperactivating all neurons within the MB, we saw a lower number of flies within 4BL (increased social space) in both males and females (**Fig. 3A**; Two-way ANOVA – Effect of neural transmission: *F_1,47_* = 7.395, *p* = 0.0091). Silencing all MB neurons showed the same reduction in the number of flies within 4BL again in both males and females (**Fig. 3A**; Two-way ANOVA – Effect of neural transmission: *F_1,30_* = 14.40, *p* = 0.0007) highlighting the importance of the MB in determining fly social space on their own. In contrast, hyperactivation of the PB neurons did not change fly social space, nor did silencing PB neurons in males but looks to have an effect in females specifically (**Fig. 3B**; Two-way ANOVA – Effect of neural transmission: *F_1,31_* = 3.443*, p* = 0.0730). Finally, we used the *nlg3-Gal4* insertion to manipulate the transmission of all *nlg3*-expressing neurons. Hyperactivation of all *nlg3*-expressing neurons decreases the number of flies within 4BL in both males and females (**Fig. 3C**; Two-way ANOVA – Effect of neural transmission: *F_1,28_* = 12.70, *p* = 0.0013). Silencing these neurons only increased social space in females but not males (**Fig. 3C**; Two-way ANOVA – Effect of interaction: *F_1,22_* = 6.169, *p* = 0.0211). There was no reduction in climbing with a hyperactivation or silencing of the MB or PB neurons (**Supp. Fig. 4A,B**), but we did observe a reduction in climbing when hyperactivating *nlg3*-expressing neurons in males only (**Supp. Fig. 4C**).

**Figure 3.**
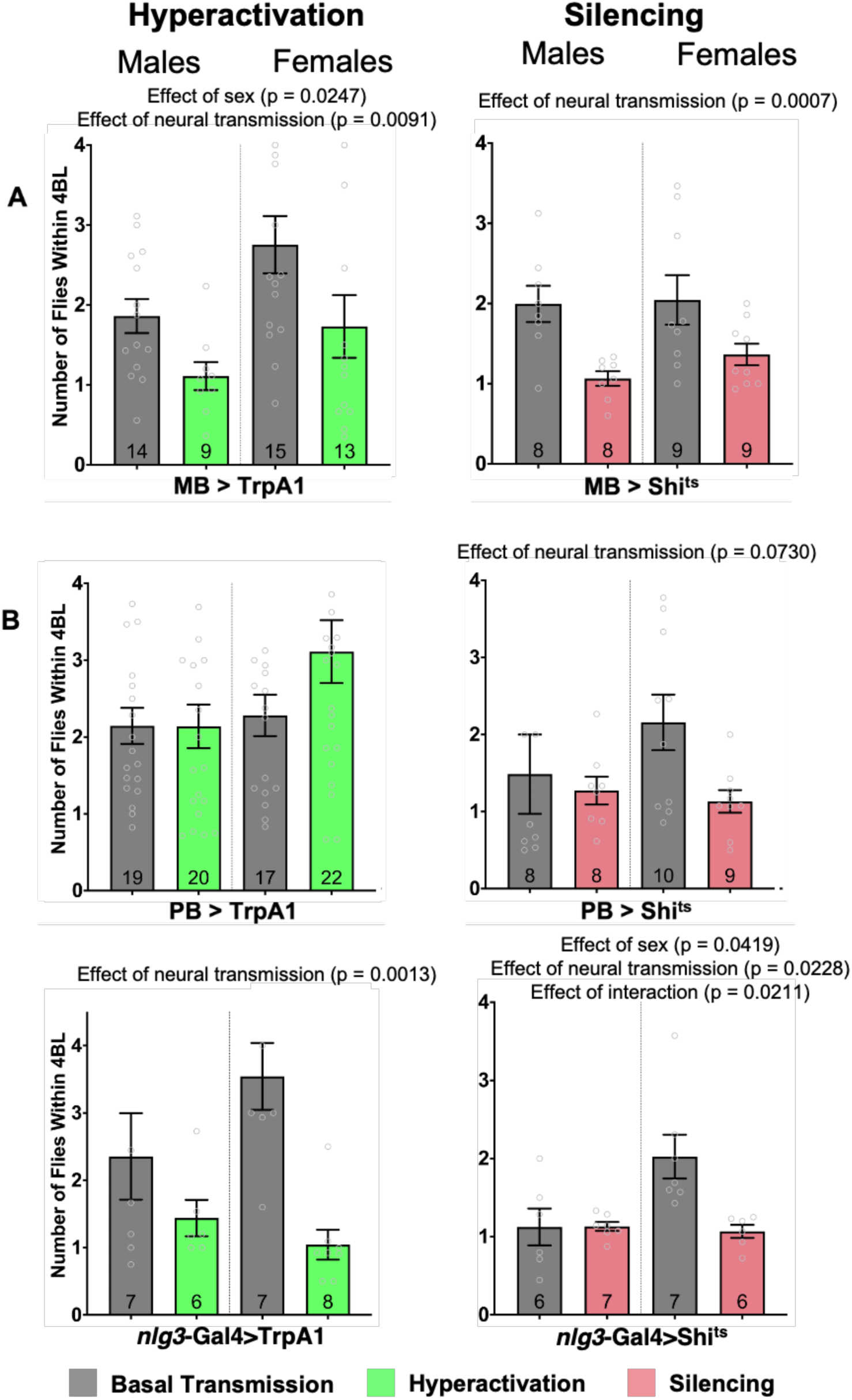
Manipulation of neural transmission within the brain structures enriched with Nlg3 increases fly social space. **A-C:** Average number of flies within 4 body lengths (4BL) in male and female flies when genetically hyperactivating or silencing neural transmission within the MB (A), PB (B), and *nlg3*-containing neurons (C). **A.** Hyperactivation of MB neurons decreases the number of flies within 4BL in both males and females compared to basal transmission of MB neurons (two-way ANOVA – Effect of neural transmission: *F_1,47_* = 7.395, *p* = 0.0091). Silencing MB neurons also decreases the number of flies within 4BL in both males and females compared to basal transmission (two-way ANOVA – Effect of neural transmission: *F_1,30_* = 14.40, *p* = 0.0007). **B.** The number of flies within 4BL was not different in males and females when the PB neurons are hyperactivated (two-way ANOVA - Effect of neural transmission: *F_1,74_* = 17.72, *p* = 0.2018) or silenced (two-way ANOVA – Effect of neural transmission: *F_1,31_* = 3.443, *p* = 0.0730). **C.** Hyperactivation of *nlg3-*containing neurons decreases the number of flies within 4BL in both males and females compared to basal transmission (two-way ANOVA – Effect of neural transmission: *F_1,28_* = 12.70, *p* = 0.0013). There is an interaction between sex and silencing *nlg3*-containing neurons where females have a decrease in the number of flies within 4BL (two-way ANOVA – Effect of interaction: *F_1,22_* = 6.169, *p* = 0.0211). Grey colour bars: inactivated genetic construct and basal transmission of neurons; green colour bars: activation of genetic construct and hyperactivation of neurons with *Gal4* driver; pink colour bars: activation of genetic construct and silencing of neurons with *Gal4* driver. The results of all two-way ANOVA and *post-hoc* tests performed can be found in **Supp. Table 2**. N = 6 – 22 for all treatments. Error bars are +/- s.e.m.

### Reducing acetylcholine release from the mushroom bodies decreases social space in both sexes, reducing GABA production only increases social space in females, but knocking down dopamine receptors in the mushroom bodies increases male social space

As the MB are important for fly social space, we investigated which neurotransmitters in the MB were important for the observed effects. Using UAS-RNAi, we knocked down the rate-limiting biosynthesis enzymes within the MB (Shih et al. 2019), including *choline acetyltransferase (ChAT* for acetylcholine production) and *glutamate decarboxylase 1* (*GAD1* for GABA production; Shih et al. 2019). Acetylcholine reduction in the MB decreased fly social space in males and females (**Fig. 4A**; Two-way ANOVA – Effect of genotype: *F_1,32_* = 11.19, *p* = 0.0021). GABA reduction also increased the number of flies within 4BL in males, but not in females (**Fig. 4B**; Two-way ANOVA – Effect of genotype: *F_1,29_* = 18.10, *p* = 0.0002).

**Figure 4.**
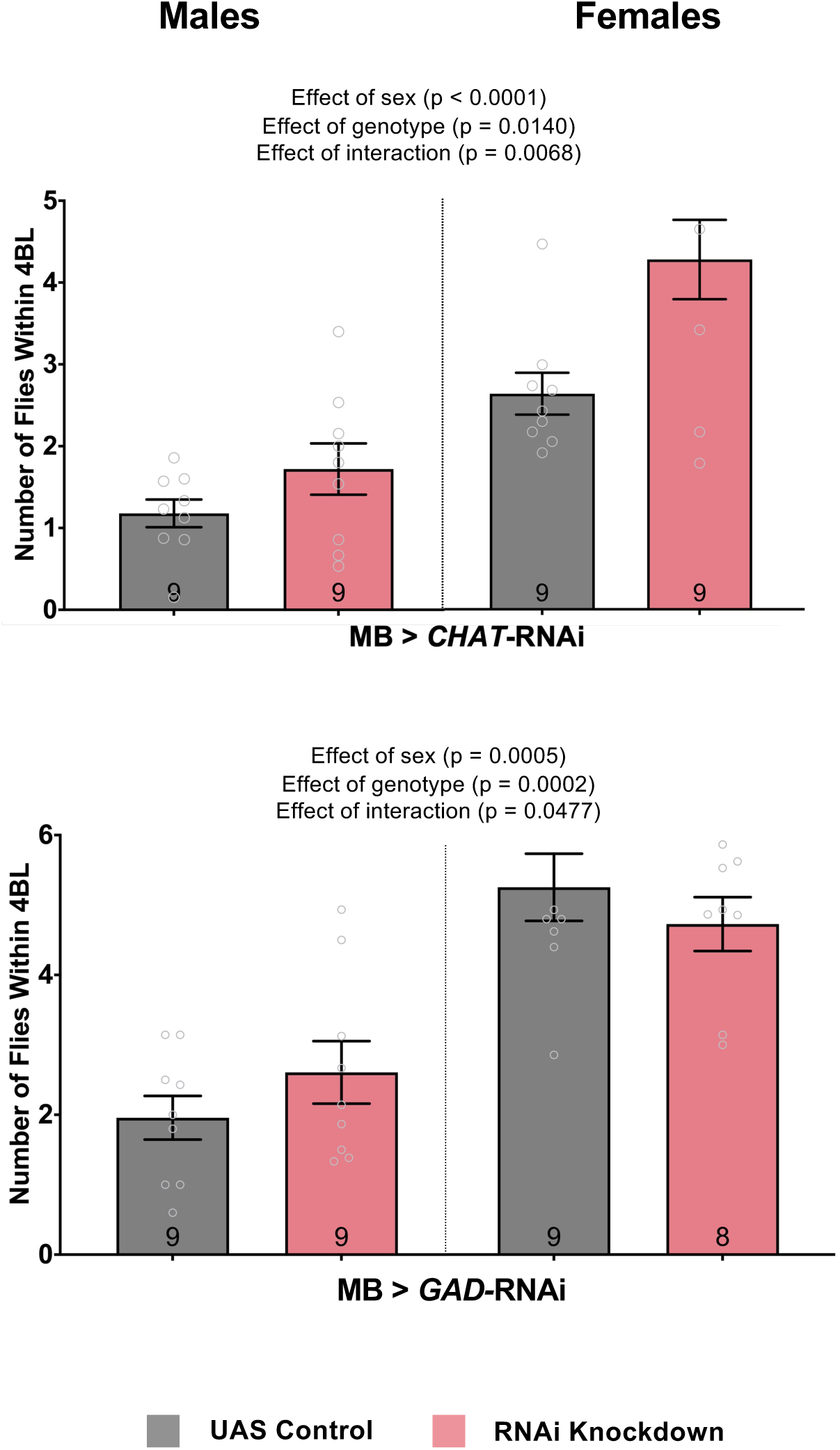
Knocking down the acetylcholine rate-limiting biosynthesis enzyme C*hAT* in the MB decreases fly social space. **A-B:** Average number of flies within 4 body lengths (4BL) in male and female flies with RNAi knockdown of *ChAT* for acetylcholine synthesis (A) or *GAD1* for GABA synthesis (B) specifically in the mushroom bodies. **A.** Both male and female flies have an increased number of flies within 4BL with *ChAT* knockdown when compared to the genetic control flies (two-way ANOVA – Effect of genotype: *F_1,32_* = 11.19, *p* = 0.0021). **B.** The interaction between sex and *GAD1* knockdown shows that male flies have an increased number of flies within 4BL while females have a decreased number compared to their genetic control flies (two-way ANOVA – Effect of interaction: *F_1,29_* = 4.276, *p* = 0.0068). Grey colour bars: genetic control *driver/+*; pink colour bars: *driver/RNAi*. The results of all two-way ANOVA and *post-hoc* tests performed can be found in **Supp. Table 2.** N = 8 – 9 for all treatments. Error bars are +/- s.e.m.

Previous research shows global dopaminergic changes in the adult fly brain will affect social space (Fernandez et al. 2017; Xie et al. 2018; Yost et al. 2020), however, the anatomical locations within the adult fly brain where alterations in dopaminergic transmission affect social behaviour is unknown. Thus, we tested the dopaminergic inputs into the MB, and found that knocking down every dopamine receptor within the MB – including Dop1R1 (**Fig. 5A**; Two-way ANOVA – Effect of genotype: *F_1,32_* = 19.61, *p* < 0.0001), Dop1R2 (**Fig. 5B**; Two-way ANOVA – Effect of genotype: *F_1,32_* = 7.401, *p* = 0.0105), Dop2R (**Fig. 5C**; Two-way ANOVA – Effect of genotype: *F_1,32_* = 41.10, *p* < 0.0001), and DopEcR (**Fig. 5D**; Two-way ANOVA – Effect of genotype: *F_1,32_* = 8.905, *p* = 0.0054) – decreased the number of flies within 4BL with a larger effect size in males than females.

**Figure 5.**
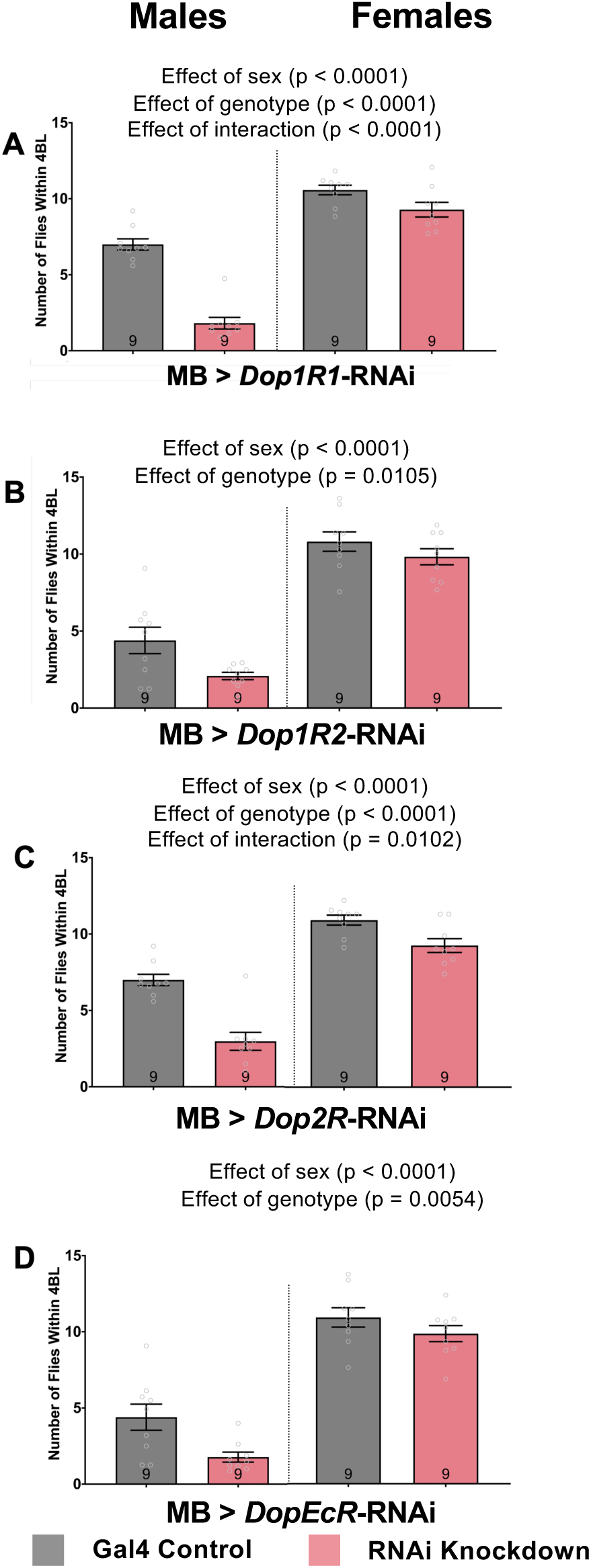
RNAi knockdown of all dopamine receptors in the MB increases social space more in males than in females. **A-D:** Average number of flies within 4 body lengths (4BL) in male and female flies knocking down dopamine receptors *Dop1R1* (A), *Dop1R2* (B), *Dop2R* (C), and *DopEcR* (D) specifically in the mushroom bodies. **A.** *Dop1R1* knockdown in the MB of males has a decreased number of flies within 4BL compared to genetic controls (Two-way ANOVA – Effect of genotype: *F_1,32_* = 19.61, *p* < 0.0001). **B.** *Dop1R2* knockdown in the MB of males has a decreased number of flies within 4BL compared to genetic controls (Two-way ANOVA – Effect of genotype: *F_1,32_* = 41.10, *p* < 0.0001). **C.** *Dop2R* knockdown in the MB of males has a decreased number of flies within 4BL compared to genetic controls (Two-way ANOVA – Effect of genotype: *F_1,32_* = 41.10, *p* < 0.0001). **D.** *DopEcR* knockdown in the MB of males has a decreased number of flies within 4BL compared to genetic controls (Two-way ANOVA – Effect of genotype: *F_1,32_* = 8.905, *p* = 0.0054). Grey colour bars: genetic control *driver/+*; pink colour bars: *driver/RNAi*. The results of all two-way ANOVA and *post-hoc* tests performed can be found in **Supp. Table 2**. N = 9 for all treatments. Error bars are +/- s.e.m.

### Neurons expressing the fruitless gene under the P1 promotor are involved in sexually dimorphic responses in fly social spacing

Since male and female flies appear to have differently neural circuitry underlying their social spacing preferences, we looked at a subset of sexually dimorphic *fruitless*-expressing neurons. We used the *fruP1-Gal4* driver to manipulate neural transmission. While hyperactivation of *fruP1* neurons decreased the number of flies within 4BL for both males and females (**Fig. 6**; Two-way ANOVA – Effect of neural transmission: *F_1,33_* = 22.65, *p* < 0.0001). silencing of *fruP1* resulted in a sex specific effect. Silencing *FruP1* neurons resulted in a higher number of flies within 4BL for males (**Fig. 6**; Two-way ANOVA and Šídák’s *post-hoc* test: *F_1,30_* = 3.451, *p* = 0.0453) but did not change female social space. Importantly, we saw a reduction in climbing in both males and females for hyperactivating (Two-way ANOVA – Effect of transmission: *F_1,32_* = 153.6, *p* < 0.0001) and silencing (Two-way ANOVA – Effect of transmission: *F_1,32_* = 33.54, *p* = 0.0102) *fruP1* neurons (**Supp. Fig. 5**).

**Figure 6.**
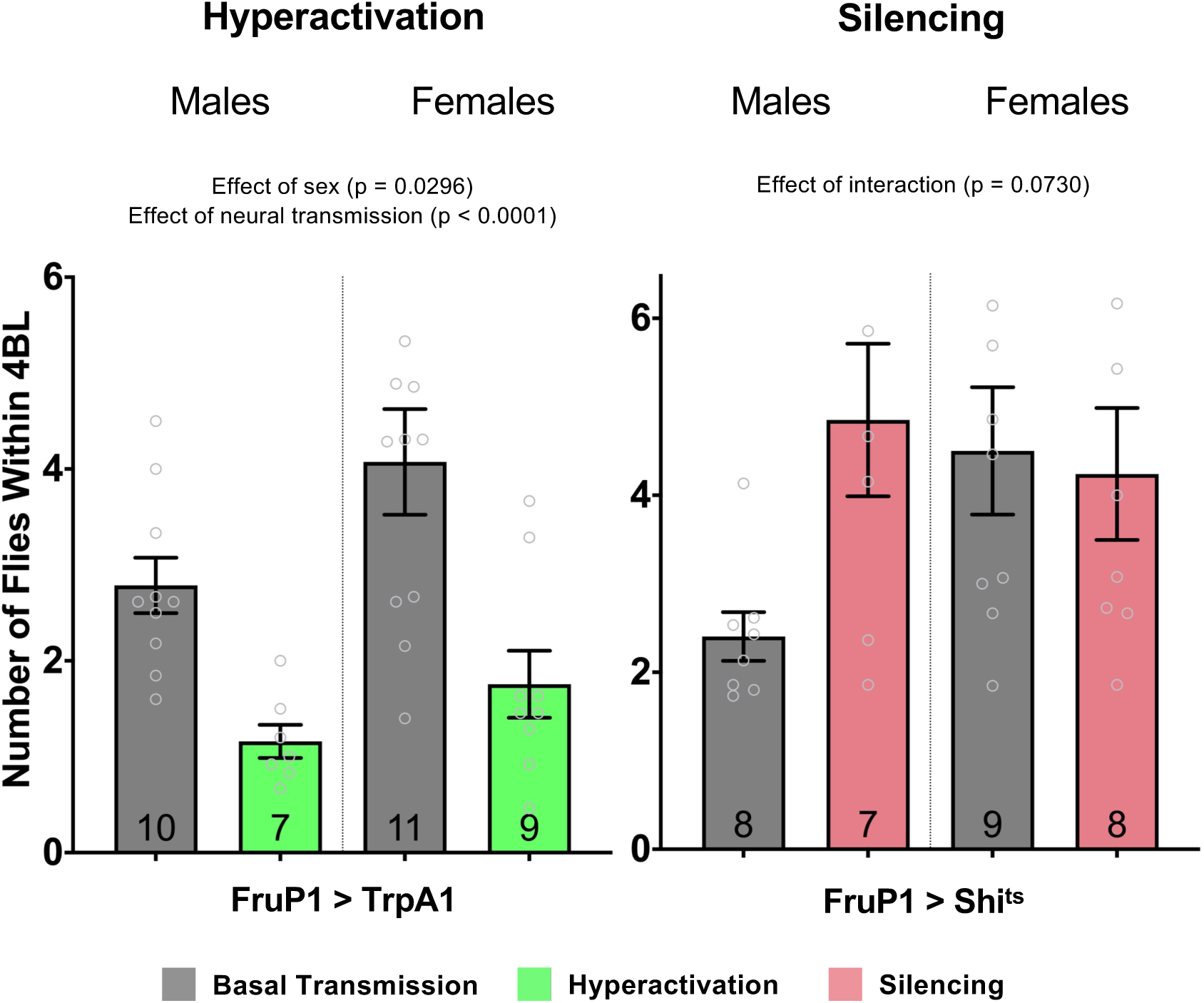
Hyperactivation of *FruP1* neurons increases social space in males and females while silencing *FruP1* neurons only decreases social space in males. Average number of flies within 4 body lengths (4BL) in males and females when genetically hyperactivating neural transmission in *FruP1* neurons. Hyperactivation of *FruP1* neurons decreases the number of flies within 4BL in both males and females compared their basal transmission (two-way ANOVA – Effect of neural transmission: *F_1,33_* = 22.65, *p* < 0.0001). Silencing *FruP1* neurons did not alter change female social space but increased the number of flies within 4BL in males (two-way ANOVA with Šídák’s *post-hoc* test: *p* = 0.0453). Grey colour bars: inactivated genetic construct and basal transmission of neurons; green colour bars: activation of genetic construct and hyperactivation of neurons with Gal4 driver; pink colour bars: activation of genetic construct and silencing of neurons with Gal4 driver. The results of all two-way ANOVA and *post-hoc* tests performed can be found in **Supp. Table 2.** N = 7 – 11 for all treatments. Error bars are +/- s.e.m.

To assess the effect of Nlg3 in *fruitless*-expressing neurons, we drove an RNAi against *nlg3* with the *fruP1* driver. We saw reducing levels of *nlg3* decreased social space in flies. Knocking down *nlg3* in *fruitless*-expressing neurons resulted in an increase in the number of flies within 4BL for males and females (**Fig. 7A**; Two-way ANOVA – Effect of genotype: *F_1,57_* = 16.27, *p* = 0.0002). Also, we observed a reduction of male climbing when knocking down *nlg3* in *fruP1* neurons (**Supp. Fig. 6A**; Two-way ANOVA – Effect of genotype: *F_1,40_* = 9.735, *p* = 0.0033). Next, we investigated the neurotransmitters involved in fly social space in *fruP1* neurons. Knocking down *TH*, the rate limiting-enzyme in the pathway of dopamine synthesis, in *fruP1* neurons did not change male or female social space (**Fig. 7B**; Two-way ANOVA – Effect of genotype: *F_1,29_* = 0.413, *p* = 0.5255). However, knocking down *TH* in *fruP1* neurons decreased climbing in both males and females indicating the importance of dopamine released from *fruP1* neurons on fly climbing (**Supp. Fig. 6B**; Two-way ANOVA – Effect of genotype: *F_1,31_* = 18.07, *p* = 0.0002). In contrast, a reduction of acetylcholine levels by knocking down *ChAT* in *fruP1* neurons increased the number of flies within 4BL in both males and females (**Fig. 7C**; Two-way ANOVA – Effect of genotype: *F_1,47_* = 99.11, *p* < 0.0001). Acetylcholine release reduction from *fruP1* neurons also decreases fly climbing (**Supp. Fig. 6C**). While a reduction of GABA by knocking down *GAD1* in *fruP1* neurons decreased male and female social space (**Fig. 8D**; Two-way ANOVA – Effect of genotype: *F_1,42_* = 8.478, *p* = 0.0057), decreasing *fruP1* GABA resulted in no change in climbing.

**Figure 7.**
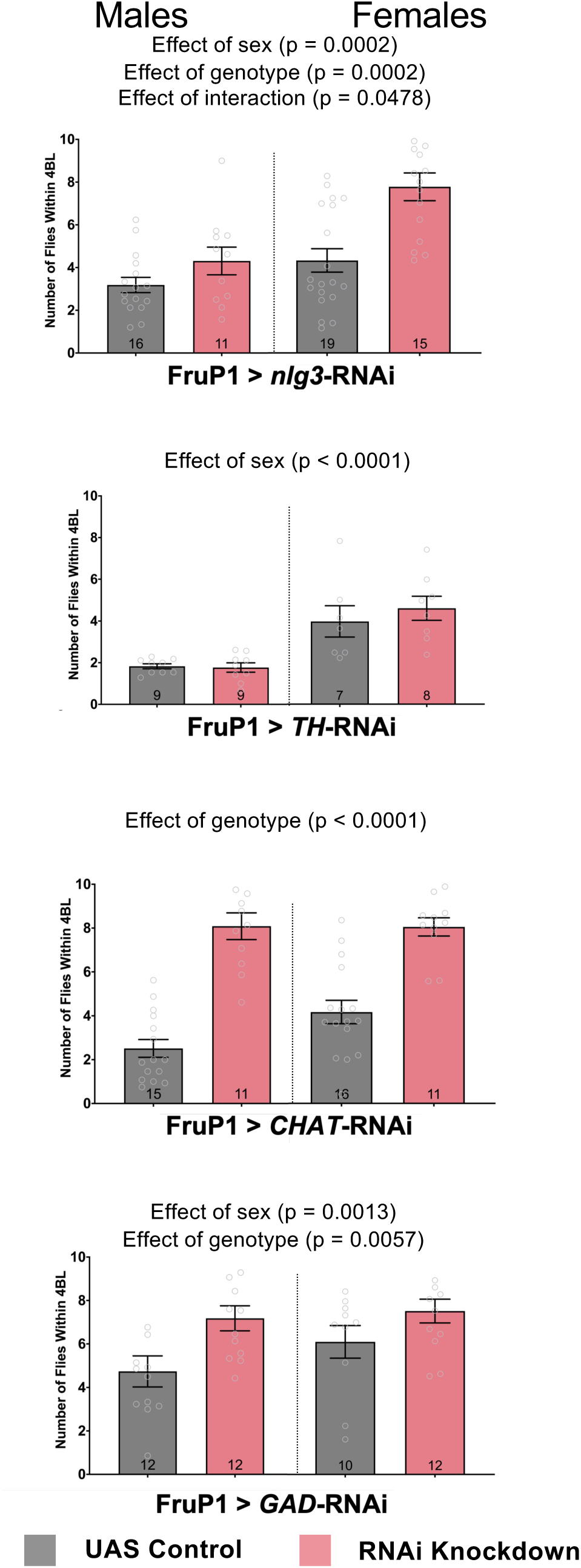
Knocking down *nlg3* in *FruP1* neurons decreases social space, while a reduction of acetylcholine and GABA in *FruP1* neurons both also decrease social space in male and female flies. **A-D**: Average number of flies within 4 body lengths (4BL) in male and female flies with RNAi knockdown of *nlg3* (A), *TH* for dopamine synthesis (B), *ChAT* for acetylcholine synthesis (C) or *GAD1* for GABA synthesis (D) specifically in *FruP1* neurons. **A.** Knocking down *nlg3* in *FruP1* has increases the number of flies within 4BL (two-way ANOVA – Effect of genotype: *F_1,57_* = 16.27, *p* = 0.0002) **B.** *TH* knockdown in *FruP1* neurons does not change social space in males and females (two-way ANOVA – Effect of genotype: *F_1,29_* = 0.413, *p* = 0.5255). **C.** *ChAT* knockdown in *FruP1* neurons increases the number of flies within 4BL in both males and females compared to the genetic control flies (two-way ANOVA – Effect of genotype: *F_1,47_* = 99.11, *p* < 0.0001). **D.** *GAD1* knockdown in *FruP1* neurons increases the number of flies within 4BL in males and females (two-way ANOVA – Effect of genotype: *F_1,42_* = 8.478, *p* = 0.0057). Grey colour bars: genetic control *driver/+*; pink colour bars: *driver/RNAi*. The results of all two-way ANOVA and *post-hoc* tests performed can be found in **Supp. Table 2**. N = 9 - 13 for all treatments. Error bars are +/- s.e.m.

**Figure 8.**
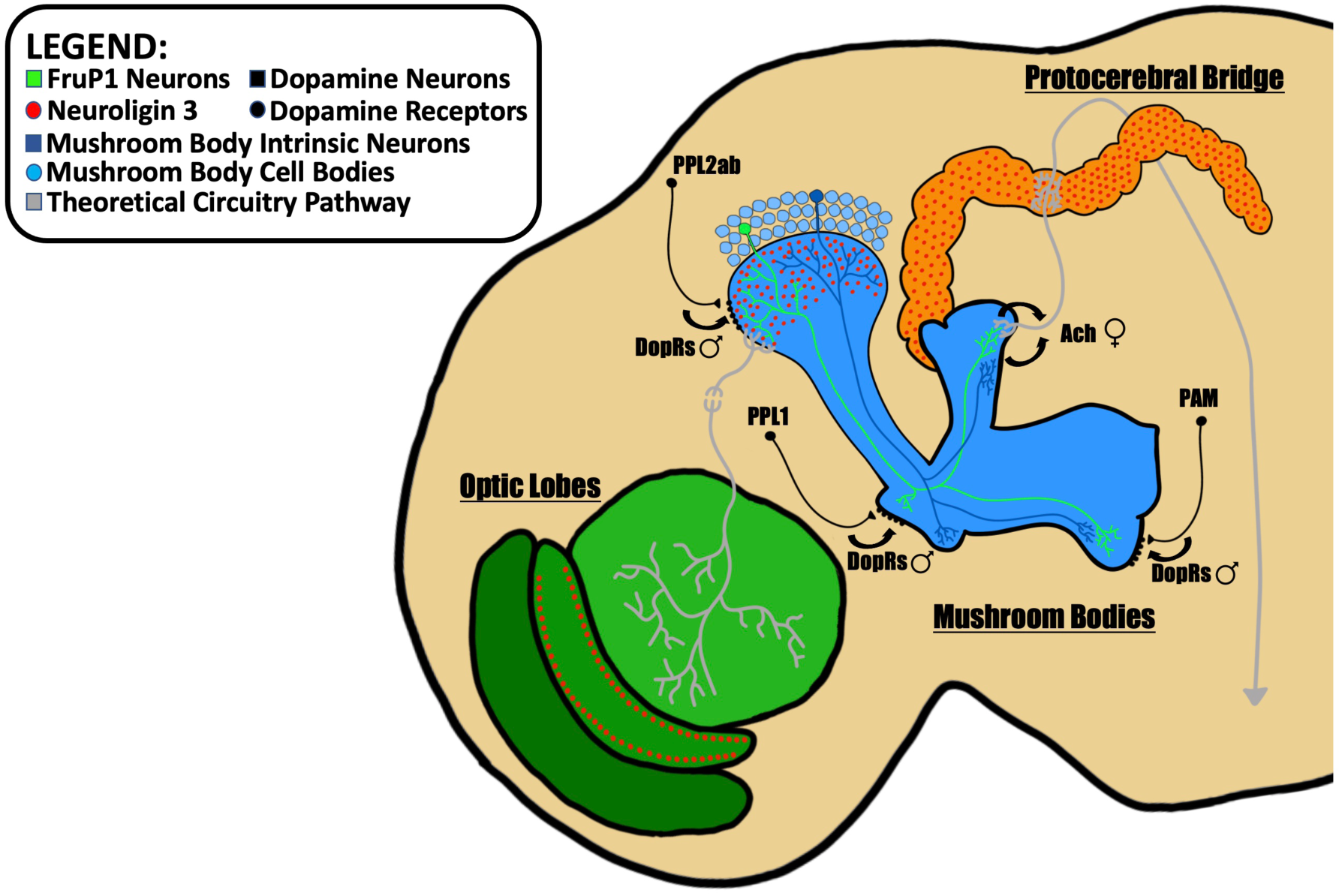
Proposed theoretical model of functional neural influences on Drosophila social space: Dopaminergic inputs into the mushroom body neurons (dark blue line) influence male social space while cholinergic outputs from the mushroom bodies influence female social space. Cholinergic outputs from *fruP1* neurons (light green line) also influence male and female social space while some mushroom body neurons also express *fru*. Neuroligin protein (red dots) enrichment is also found to be located in the optic lobes, calyx of the mushroom bodies, and protocerebral bridge.

## Discussion

Social interactions within a group requires an integration of both attractive and repulsive social cues that lead to a behavioural output. Here, we expand the map of the neural basis for social behaviour by focusing on the sexually dimorphic role of *neuroligin 3* and its associated adult brain structures in male and female *Drosophila melanogaster*.

We show for the first time that *nlg3* is expressed at low levels in the mushroom bodies, optic lobes, and protocerebral bridge in the adult fly brain. Each of those structures independently requires *nlg3* for normal social spacing. The importance of Nlg3 in the OL on social behaviour was expected as vision is known to be important for proper social spacing (Simon et al. 2012; Burg et al. 2013; Jiang et al. 2013).

Additionally, manipulating total neural transmission in these brain regions highlighted the importance of the Kenyon cells for maintaining social space. Hyperactivating and silencing MB neurons in both males and females increased fly social space. However, hyperactivating and silencing the PB only affected social space in females. Reducing acetylcholine release from the MB decreased social space in both sexes, while reducing GABA only decreased social space in males and knocking down all dopamine-type receptors increased mostly male social space. These findings suggest that mapping the circuitry through the MB happens in a sexually dimorphic manner, where total acetylcholine release from the MB will modify female social space and dopaminergic inputs to the MB will change male social space. We still do not know the specific output from the MB leading to changes in males or the inputs to the MB leading to changes in females.

We also examined the involvement of *fru* neurons, which are known to regulate other sexually dimorphic behaviors such as male courtship or female receptivity (Chowdhury et al. 2020). Hyperactivating *fruP1* neurons decreased social space in both sexes, while silencing them showed to be sexually dimorphic with increased social space only occurring in males. Again, this demonstrates the influence *fruP1* neurons have on social space along with other fly social behaviours.

Lastly, manipulation of neurotransmitters within *fruP1* neurons revealed that the release of dopamine form those neurons was not important for social space. Instead, acetylcholine release from *fruP1* neurons play a role in fly social space, a phenocopy observed when knocking down acetylcholine in the MB, but GABA release decreased social space in both, with a smaller effect size in females. In addition, RNAi knockdown of *nlg3* in the *fruP1* neurons also showed sex-specific effects on social space, with a larger effect size in females.

Our antibody targeted the intracellular domain of Nlg3, which allowed us to identify protein localization specifically in the calyx (dendritic ends of the Kenyon cells) of the mushroom bodies, the medulla and lamina layers of the optic lobes, and throughout the entire protocerebral bridge (**Fig. 1**). However, there was a low signal to noise ratio when imaging with just a polyclonal anti-Nlg3 antibody (**Fig. 1A,B**) indicating a low level of expression of *nlg3* in the adult 3-4 day old brain. Attempts to create another polyclonal, or even a monoclonal antibody specific to Nlg3 were unsuccessful, so we transitioned to using two different genetic constructs (*MiMIC-nlg3* and *nlg3-Gal4 > mCD8-GFP*) to confirm the results we observed with our polyclonal antibody. The GFP localization in this line in the mushroom bodies and optic lobes was similar to the Nlg3 protein localization (**Fig. 1C,D**). However, the GFP expression in the *MiMIC-nlg3* was also low, probably due to the endogenously low expression of *nlg3* (reported in Yost et al. 2020), and we could not see any expression in the protocerebral bridge. We then switched to a *nlg3-Gal4* driver to express Gal4 transcription factor in the same cells as *nlg3*, allowing to activated the *UAS-GFP* at higher levels than just endogenous *nlg3* (**Fig. 1F**) allowing for better visualization with immunofluorescence of the calyx and protocerebral bridge (**Fig. 1E**).

After determining the localization of *nlg3* expression in the adult brain, we examined the effects of targeted RNAi knockdown of *nlg3* on fly social space. Deletions of the whole *nlg3* gene leads to an increase in fly social space (Yost et al. 2020), which we replicated with an RNAi knockdown. Knocking down *nlg3* in the optic lobes might affect fly vision but we did not yet directly test this. The RNAi knockdown of *nlg3* leads to a reduction in climbing (**Supp. Fig. 3C**) as the climbing assay is performed under uniform light. The reduction in climbing may be influenced by impairments in vision and not solely or entirely due to impairments in locomotion. Vision is known to play a role in social spacing as well (Simon et al. 2012; Burg et al. 2013; Jiang et al. 2020), so the role of *nlg3* on vision will need to be further investigated in the future.

Notably, we observed that the knockdown of *nlg3* in all three structures individually resulted in sex differences in social space with females generally settling in a group closer together than males (**Fig. 2A-C**). This is in line with previously published results with sex differences seen in social space (Simon et al. 2012).

We then modified the transmission of the neurons within these structures to assess their individual roles in this social behaviour. Indeed, Burg et al. (2012) reported the importance of the MB in fly social space. We also observed an increase to male and female social space whether the entire MB neurons were hyperactivated or silenced (**Fig. 3A**). As the MB is not only important for learning and memory but also sensory integration, it would follow that manipulation of this circuitry would affect behavioural output.

There were no changes to social space when we hyperactivated the PB neurons but silencing the PB neurons increased female social space specifically (**Fig. 3B**) suggesting the important of the PB circuitry in regulating female social space. Next, using the *nlg3-Gal4* driver, we manipulated the transmission of all *nlg3*-expressing neurons revealing their sexually dimorphic effect on social space. Females will get further apart with hyperactivation or silencing of *nlg3*-containing neurons, but males will only get further apart with hyperactivation and not silencing of the same neurons (**Fig. 3C**). This might explain the sexually dimorphic effect of *nlg3* deletion on flies of the same age (Yost et al. 2020) are not influenced by just one of the identified brain structures *nlg3* is expressed in (MB, PB, OL), but instead may be due to the collective circuitry of all *nlg3*-expression neurons (**Fig. 1**).

The mushroom bodies influence on fly social space was clear with changes to neural transmission, so we then aimed to identify the neurotransmitter released from the mushroom bodies that affected social spacing. As expected, knocking down *ChAT* and reducing acetylcholine levels in the MB decreased social space in both sexes (**Fig. 4A**). But knocking down *GAD* and reducing GABA levels in the MB decreased social space only in males (**Fig. 4A**).

We also found that knocking down all dopamine receptor types in the mushroom bodies will specifically increase male social space but did not affect females (**Fig. 5A-D**). Of note, Dop1R1 and Dop2R knockdown did increase female social space (**Fig. 5A,C**; Two-way ANOVA – Effect of interaction: *F_1,32_* = 27.24, *p* < 0.0001; Two-way ANOVA – Effect of interaction: *F_1,32_* = 7.448, *p* < 0.0001, respectively) but this was only a small difference in the number of flies within 4BL and was at the upper end of how close flies can group together showing a ceiling effect that produced a minimal change in their behaviour. Overall, this would indicate that the transmission of the Kenyon cells themselves, specifically acetylcholine release, will influence female social space but the dopaminergic inputs into the MB modify the male decision of their social space. It was interesting to observe that all the 4 types of Drosophila dopamine receptors (Dop1R1, Dop1R2, Dop2R, and DopEcR) all had the same effect on social space while each receptor plays unique roles to fly behaviour such as sleep, courtship, feeding, and memory (Kim et al. 2003; Lebestky et al. 2009; Kuo et al. 2015).

In Drosophila social space, many genetic and neural manipulations lead to sex differences, even though both sexes respond to changes in their social environment in similar ways. For example, age, mating, and isolation all lead to decreased social space in both sexes (Simon et al. 2012; Brenman-Suttner et al. 2018; Yost et al. 2024). Due to this, we examined the influence that the sexually dimorphic *fruP1* neuron subset had on social space. Again, we saw male-female sex differences in response to modulation of *fruP1* neuronal transmission. For males, when *fruP1* neurons are hyperactivated, flies get further apart, but when these neurons are silenced, males get closer together (**Fig. 6**). Females also get further apart when *fruP1* neurons are hyperactivated, but do not change their social space when these neurons are silenced (**Fig. 6**). This indicates that *fruP1* neurons in males can increase and decrease neuronal activity resulting in a change in their social space, but for females only an increase in *fruP1* activity modifies their social space. The basal transmission of these neurons that affect female social space might not be active and therefore cannot be silenced, only hyperactivation changes this resting state and leads to females settling further in a group setting.

Knowing the importance for *fruP1* neurons in Drosophila social space, we then examined the molecular players within these neurons that are contributing to the behavioural changes. Knocking down *nlg3* in *fruP1* neurons showed a male-female sexual dimorphism with males getting further apart and females closer together (**Fig. 7A**). This confirms there are *fruitless-* and *nlg3*-expressing neurons that are contributing to fly social space behaviour. Furthermore, our results indicate that those neurons are not dopaminergic. However, the *fruP1* neurons affecting social space are cholinergic, in both males and females, a phenocopy of the effect seen on females both when reducing acetylcholine levels in the mushroom bodies and when knocking down *nlg3* in *fruP1* neurons. Because a subset of the MB neurons is also *fruP1* neurons (Palmateer et al. 2023), we propose that the Kenyon cells *nlg3* neurons are also *fruP1* neurons, at least those that affect female social space (**Fig. 8**).

Finally, as previously reported (Jiang et al. 2020; Yost et al. 2024), we did not observe any correlation in our experiments between variations in performance index leading to any changes in social space.

Fly social behaviour relies heavily on an individual’s ability to perceive and integrate social and environmental cues. Using the Drosophila social space assay, we show the roles of the OL and MB in regulating fly social spacing. The OL are essential for sensory perception, as they transduce visual cues into electrochemical signals that are integrated within MB, and the central complex, thereby enabling the fly to perceive its environment and alter its behaviour accordingly.

The significance of Nlg3 in the OL is reiterated here, as it is probably involved in the proper development and regulation of synaptic connections within these synaptically rich brain regions. Proper development of the OL layers is required to decode parallel circuitry pathways involved in processing color, shapes, and motion, which are crucial for interpreting environmental stimuli. Outside of the development and maintenance of synaptic connections that can affect fly behaviour, neural transmission through these brain regions reveals the MB to be important for social behaviour. The MB, which receives signals and integrates from multiple other brain regions (Aso et al. 2014), relies on proper development of synaptic connections where disruptions or improper regulations in these connections can lead to misinterpretation of signals, ultimately altering fly behavior through its descending output neurons.

By dissecting the molecular players within these neuronal subsets, we identify some of the inputs and outputs of the MB that influence fly social spacing. This behavioral data allows us to begin constructing a functional circuitry model of the neural pathways that regulate social space and where sexual dimorphic integration of cues might be taking place (hypothesis generating model **Fig. 8**). We have identified that dopaminergic signalling into the MB influences social space but still need to identify which dopaminergic clusters are playing a role. We can similarly test the known dopaminergic subsets to innervate the MB (PPL1, PAM, and PPL2ab). We also identified that acetylcholine signalling from the MB will affect female social space, but we have yet to identify which mushroom body output neurons (MBONs) are associated with these behavioural changes (**Fig. 8**). We can also test the known 21 MBONs to find downstream circuits that influence this social behaviour specifically.

## Acknowledgments

We want to thank Dr. Michael Strong in whose laboratory the immunocytochemistry imaging work was performed, and for the help of his research team for the troubleshooting involved in obtaining good quality images.

This study would not have been possible without the continued funding received by Western’s Biotron Facility and the support offered there by the staff: we are very thankful to Karen Nygard for her technical and molecular help in immunofluorescence protocol establishment and microscopy within the integrated microscopy unit. We also want to thank the Insect Suite within the Biotron Facility of Western University, for housing our flies in climate-controlled walk-in rooms (25°C). We also acknowledge Dr. Ryley T. Yost for his western blot analysis to show the *nlg3-RNAi* results in a reduction of Nlg3 protein levels (data not shown).

We also want to thank Flybase (Öztürk-Çolak et al., 2024 - NHGRI Awards U41HG000739 and U24HG010859), the Bloomington Drosophila Stock Center (NIH P40OD018537) and for the nc82 (anti-Brp) primary antibody which was obtained from the Developmental Studies Hybridoma Bank, created by the NICHD of the NIH and maintained at The University of Iowa, Department of Biology, Iowa City, IA 52242, for the invaluable resources they provide to the scientific community.

## Ethics Statement

Western’s Animal Care Committee or Ontario Provincial and Federal regulatory bodies specifies that ethical considerations do not apply to the study of invertebrate animals including insects such as *Drosophila melanogaster*

## Data availability statement

The datasets analysed for this study will be deposited in the Dryad repository (TBD).

## Authors contribution

Based on Oxford Academic (Genetics journal)

JWR: Conceptualization, Supervision, Project Administration, Investigation, Formal Analysis, Methodology, Resources, Funding Acquisition, Writing - Original Draft Preparation, and Writing - Review & Editing.

ATB, MRE: Conceptualization, Investigation, Formal Analysis, Funding Acquisition, and Writing - Review & Editing.

JK, AR: Conceptualization, Investigation, Formal Analysis and Writing.

JRI: Methodology, Review & Editing.

SP, DSL: Investigation and Formal Analysis.

AS: Methodology.

JND: Investigation, Formal Analysis and Validation.

GIR: Writing - Review & Editing.

AJM: Methodology, Supervision, and Funding Acquisition.

AFS: Conceptualization, Supervision, Project Administration, Formal Analysis, Resources, Funding Acquisition, and Writing - Review & Editing.

## Supplemental Figure Captions

**Supplemental Figure 1:**
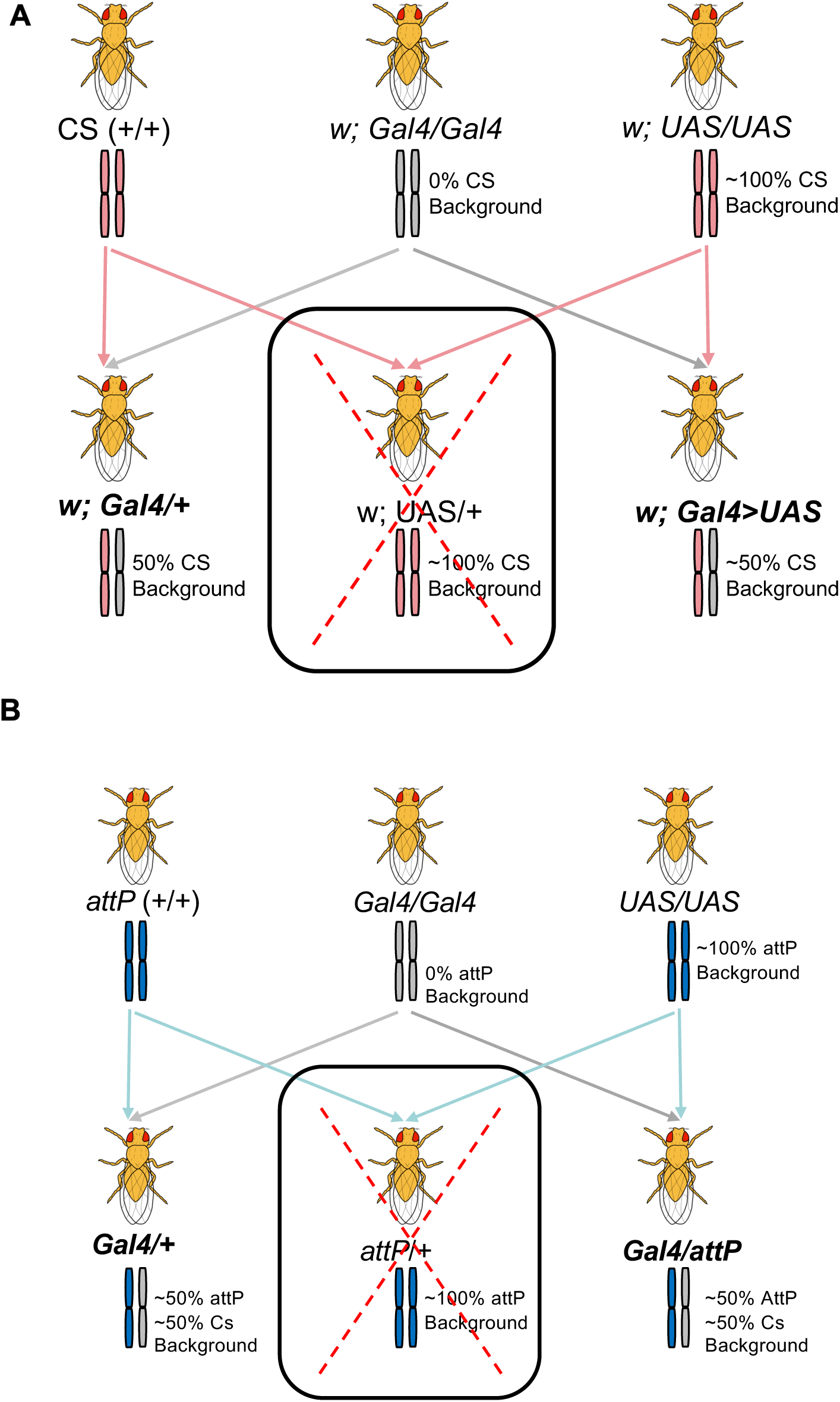
Schematic of crossing scheme used to generate genotype controls for all RNAi knockdown experiments. Bolded genotypes are presenting in main figures shown. Results for both genetic controls are included in **Supplemental** Figures 7-10. (A) Genetic crossing scheme used for Cs background and exclusion of inappropriate genetic background as a control. (B) Genetic crossing scheme used for V10 background and exclusion of inappropriate genetic background as a control.

**Supplemental Figure 2:**
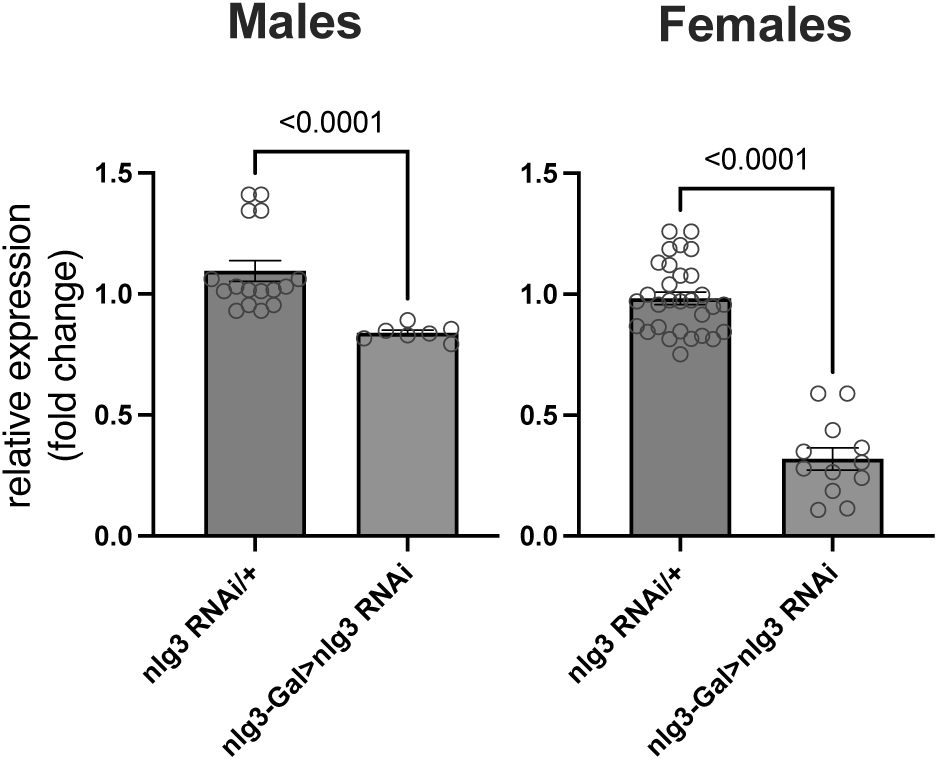
Relative expression of *nlg3* mRNA in male (M) and female (F) fly heads (3–5 days old) was measured by qPCR. Data represent mean fold-change (± SEM) from 3 biological replicates with 3 technical replicates each, with Tubulin84 as the reference gene; statistical evidence was assessed using a unpaired t-test.

**Supplemental Figure 3:**
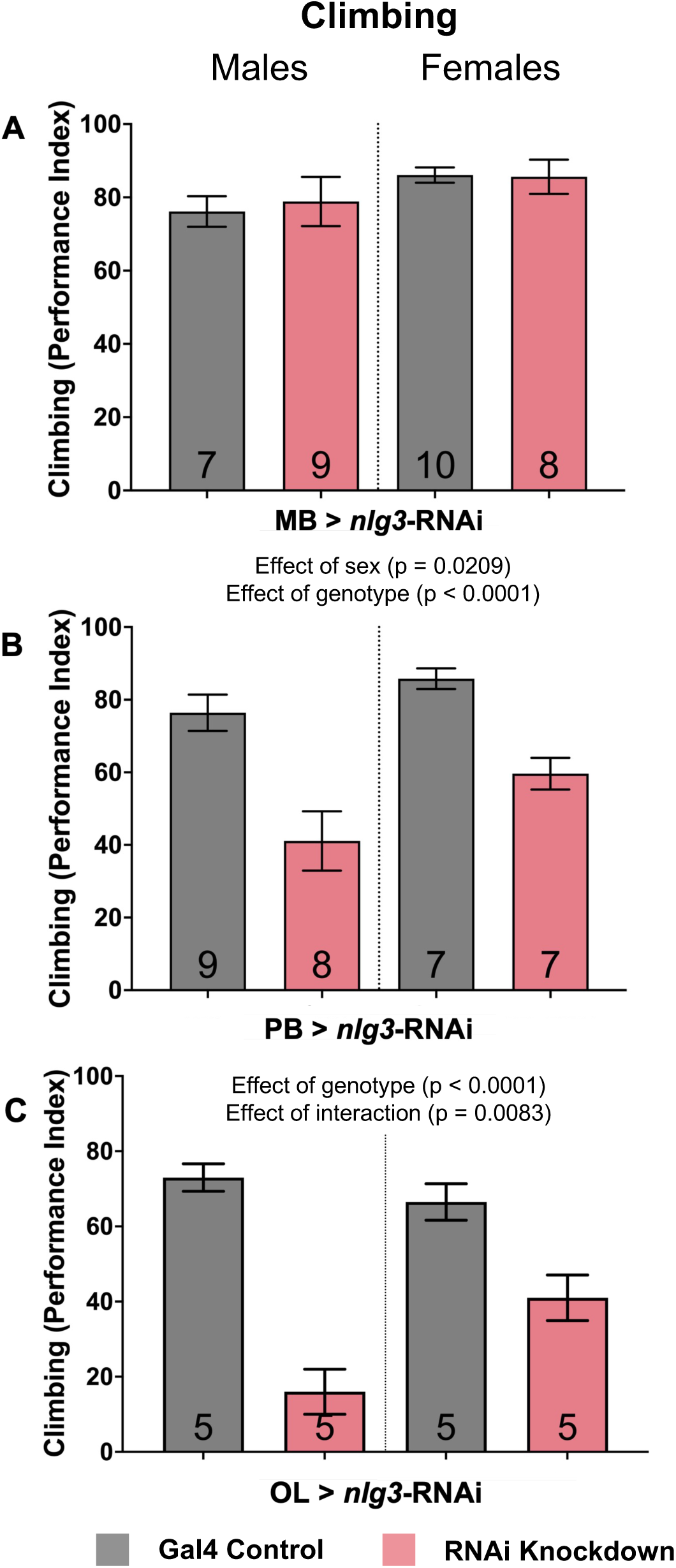
Knocking down Nlg3 specifically in the Protocerebral bridge and optic lobes decreases fly climbing in males and females but not when knocking down in the mushroom bodies specifically. **A-C:** Startle-induced climbing represented by the mean percentage of flies reaching the top vial (performance index). Genetic constructs include RNAi knockdown of *nlg3* targeted specifically to the mushroom bodies (A), protocerebral bridge (B), and the optic lobes (C). Grey colour: genetic control *driver/+*; pink colour: *driver/RNAi*. The results of all two-way ANOVA performed are shown on the figure. N = 5 – 10 for all treatments. Errors bar are +/- s.e.m.

**Supplemental Figure 4:**
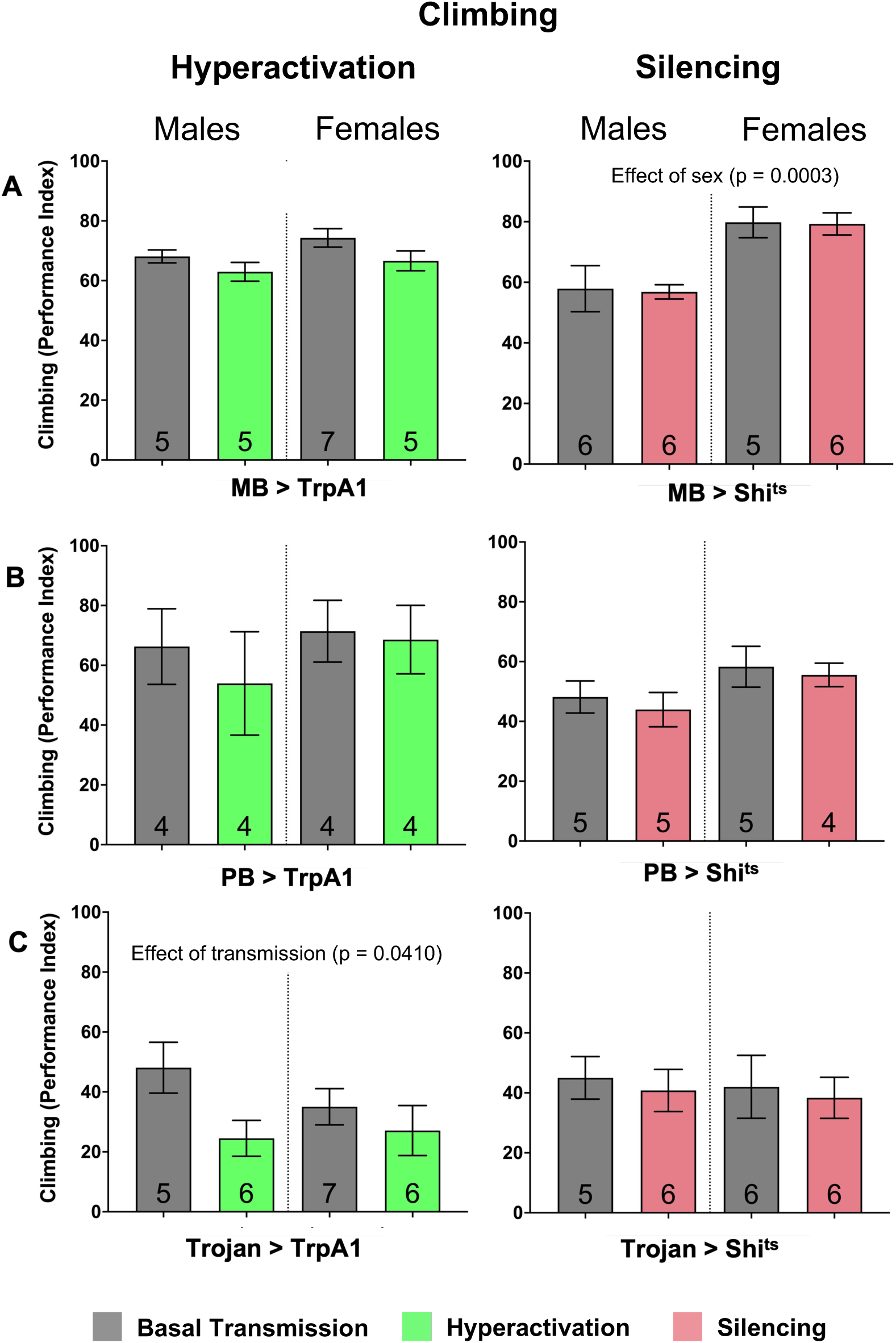
Hyperactivation of all the brain structures enriched with Nlg3 decreases fly climbing. **A-C:** Startle-induced climbing represented by the mean percentage of flies reaching the top vial (performance index). Hyperactivation or silencing neurons specifically to the mushroom bodies (A), protocerebral bridge (B), and all *nlg3-expressing* neurons (C). Grey colour bars: inactivated genetic construct and basal transmission of neurons; green colour bars: activation of genetic construct and hyperactivation of neurons with *Gal4* driver; pink colour bars: activation of genetic construct and silencing of neurons with *Gal4* driver. The results of all two-way ANOVA performed are shown on the figure. N = 4 – 7 for all treatments. Error bars are +/- s.e.m.

**Supplemental Figure 5:**
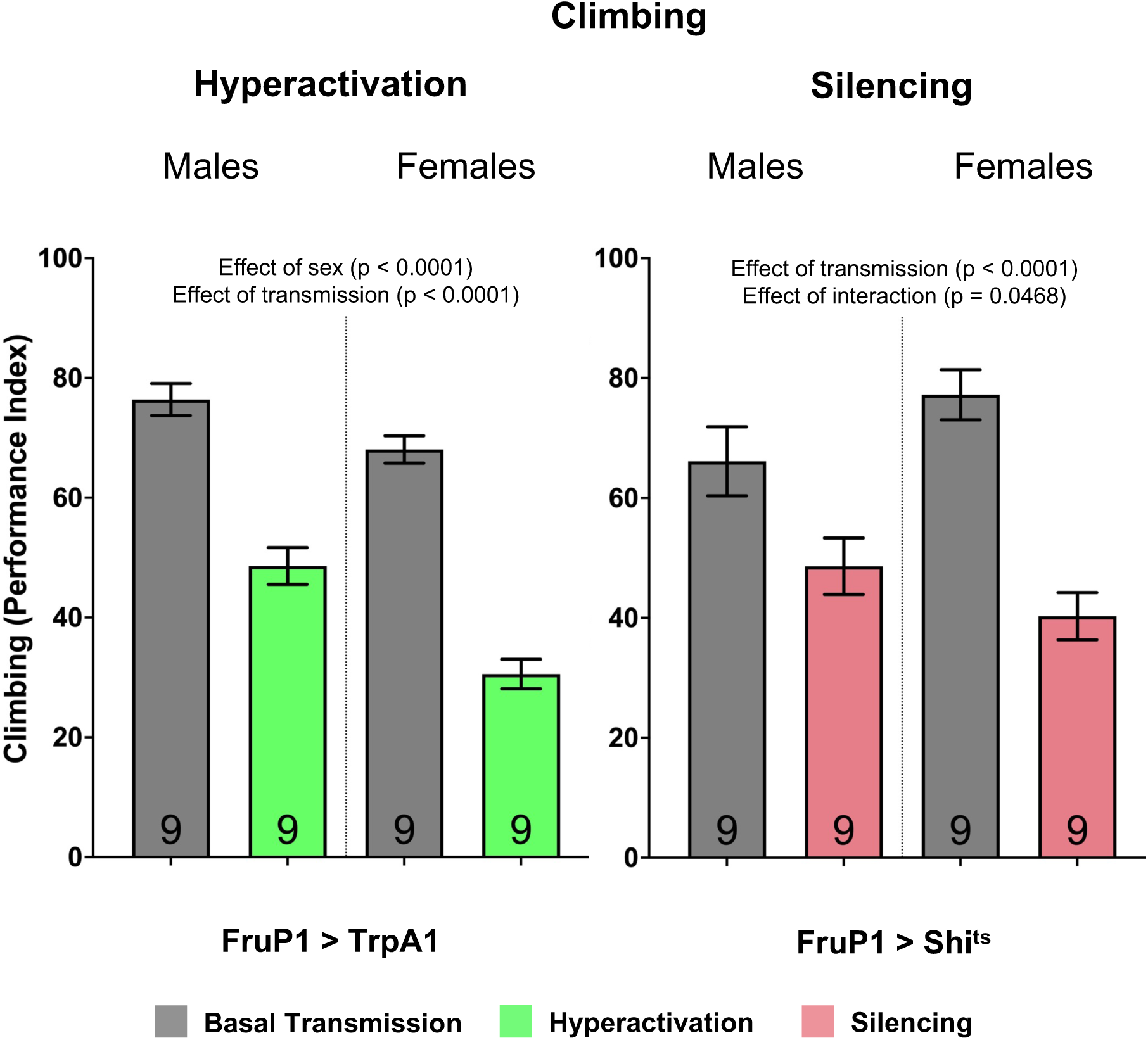
Hyperactivation and silencing of *FruP1* neurons decreases climbing in males and females. Startle-induced climbing represented by the mean percentage of flies reaching the top vial (performance index). Hyperactivation or silencing neurons specifically in *FruP1* neuron subset. Grey colour bars: inactivated genetic construct and basal transmission of neurons; green colour bars: activation of genetic construct and hyperactivation of neurons with *Gal4* driver; pink colour bars: activation of genetic construct and silencing of neurons with *Gal4* driver. The results of all two-way ANOVA performed are shown on the figure. N = 9 for all treatments. Error bars are +/- s.e.m.

**Supplemental Figure 6:**
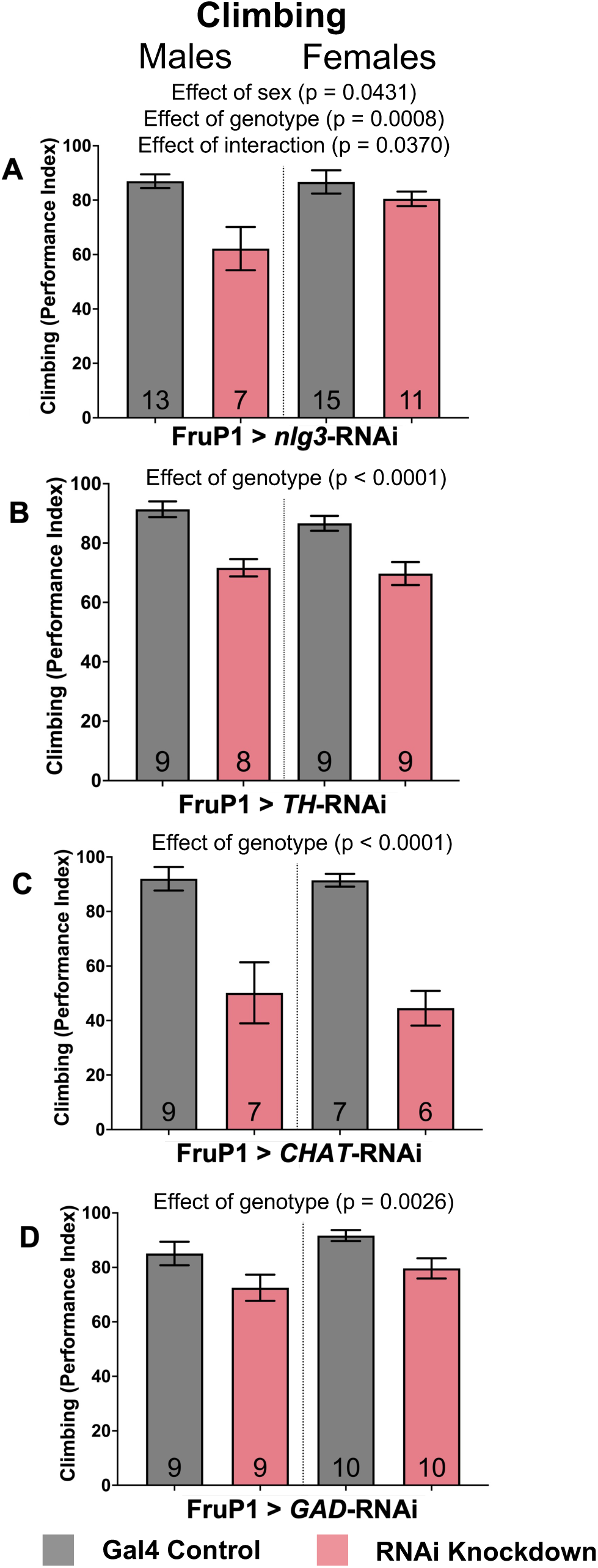
Knocking down *nlg3* in *FruP1* neurons decreases male climbing, while a reduction of dopamine, acetylcholine, and GABA in *FruP1* neurons all decrease climbing in male and female flies. **A-D:** Startle-induced climbing represented by the mean percentage of flies reaching the top vial (performance index). Genetic constructs include RNAi knockdown targeted specifically to the *FruP1* subset of neurons of *nlg3* (A), knocking down dopamine (B), acetylcholine (C), and GABA (D). Grey colour: genetic control *driver/+*; pink colour: *driver/RNAi*. The results of all two-way ANOVA performed are shown on the figure. N = 6 – 15 for all treatments. Errors bar are +/- s.e.m.

**Supplemental Figure 7:**
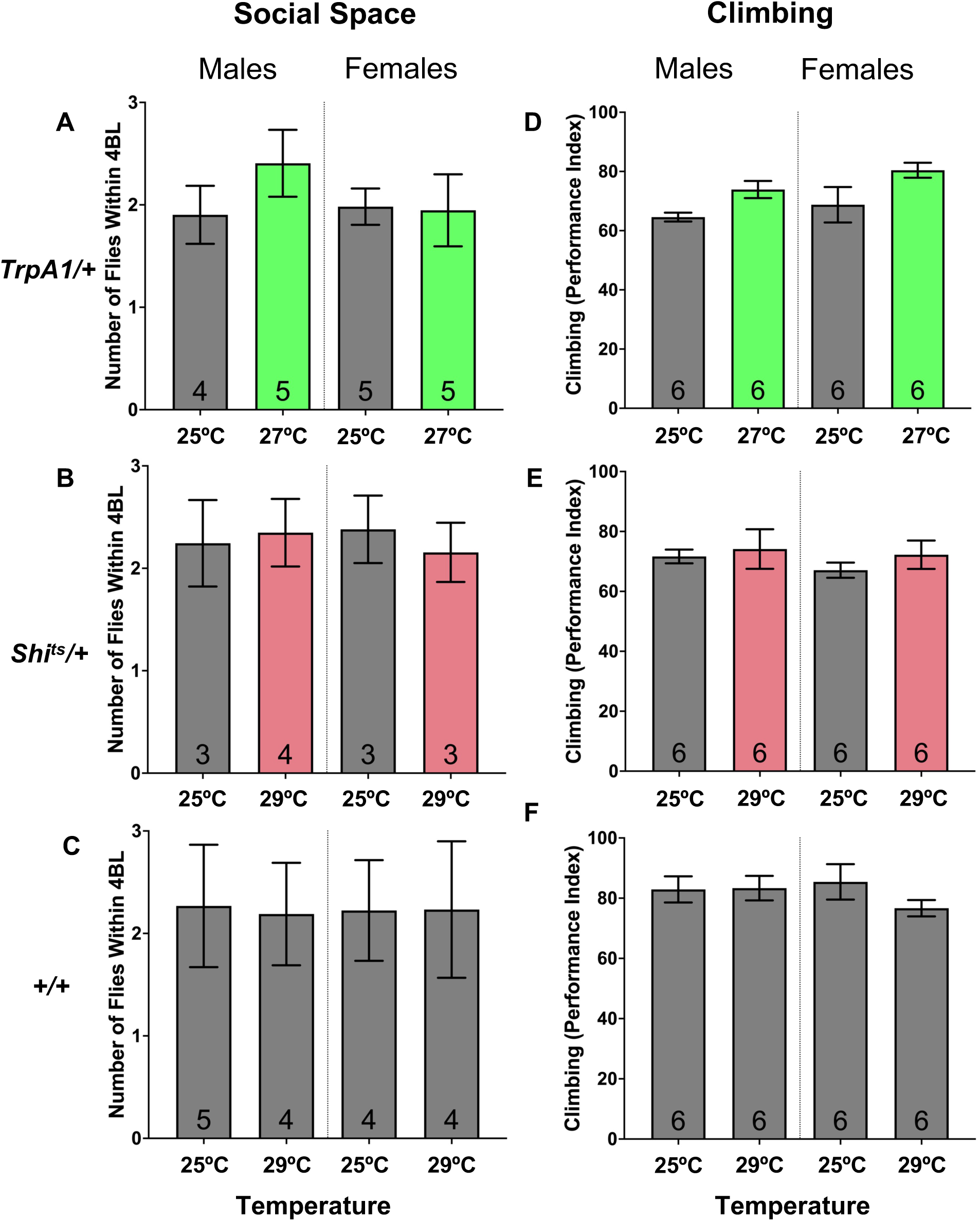
Elevated temperature on Cs, *Trp1A/+*, and *Shi^ts^/+* does not affect social space or climbing. The genetic controls of the temperature sensitive constructs were tested in social space with the average number of flies within 4 body lengths **A-C:** and climbing with a performance of percentage of flies reaching the top vial **D-F:** in both male and female which reveals the construct alone does not affect social space or climbing.

**Supplemental Figure 8:**
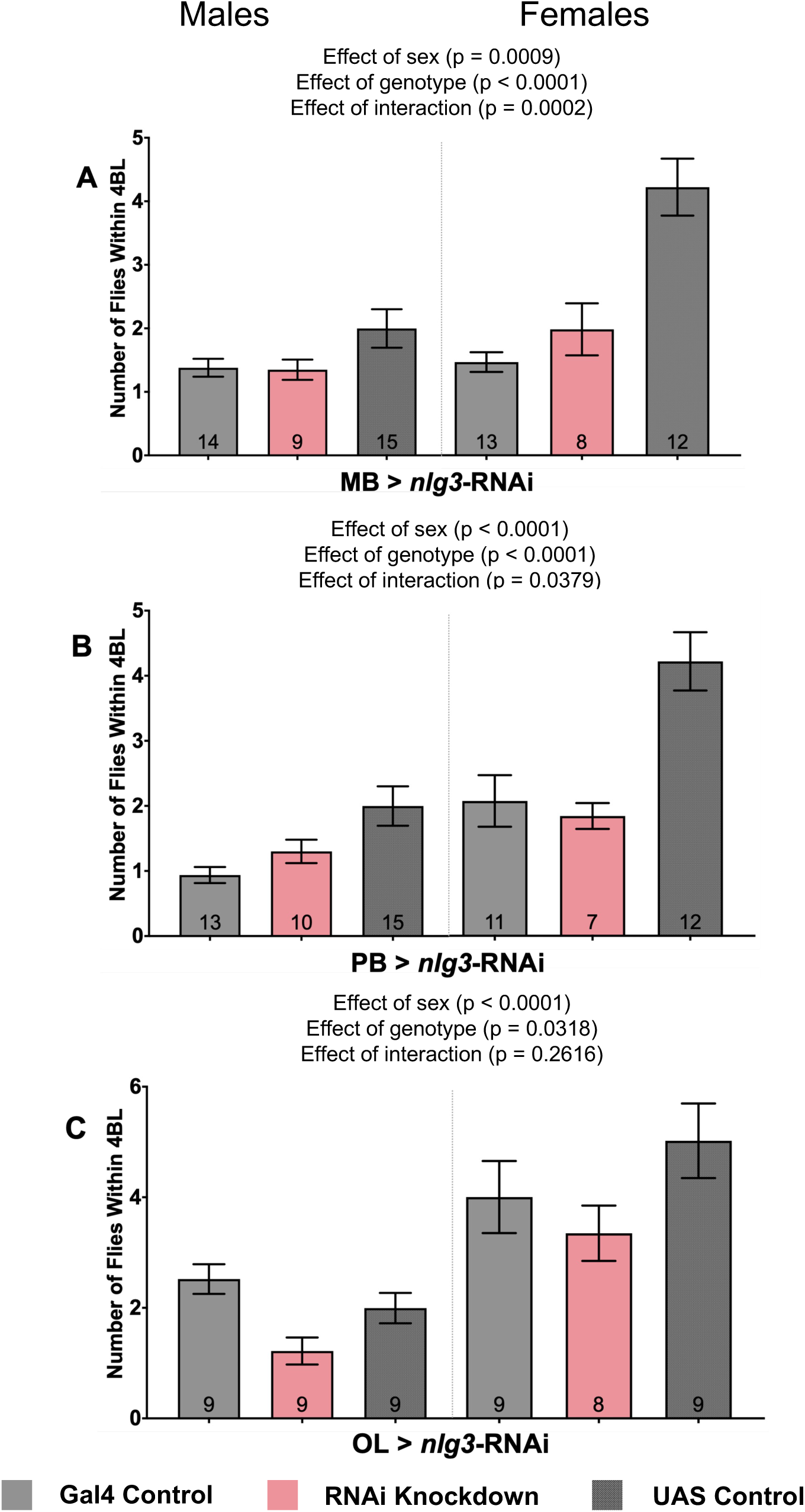
Knocking down Nlg3 specifically in the optic lobes increases fly social space in males and females but no changes are seen when knocking down in the protocerebral bridge or mushroom bodies specifically when looking at both genetic controls with different genetic backgrounds. Average number of flies within 4 body lengths (4BL) in male and female flies with RNAi knockdown of *nlg3* targeted specifically to the mushroom bodies (A), protocerebral bridge (B), and the optic lobes (C). **A.** The number of flies within 4BL does not change in both males and females in flies with *nlg3* knockdown targeted to the MB. Comparing the knockdown line to both control lines shows an intermediate phenotype. **B.** The average number of flies within 4BL does not change in both males and females in flies with *nlg3* knockdown targeted to the PB. Comparing the knockdown line to both control lines shows an intermediate phenotype. **C.** Knocking down *nlg3* in the OL decreases the number of flies within 4BL in both males and females compared to genetic control flies (two-way ANOVA – Effect of genotype: *p* = 0.0318). Grey colour: genetic control *driver/+* and *Gal4/+*; pink colour: *driver/RNAi*. N = 7 – 14 for all treatments. Error bars are +/- s.e.m.

**Supplemental Figure 9.**
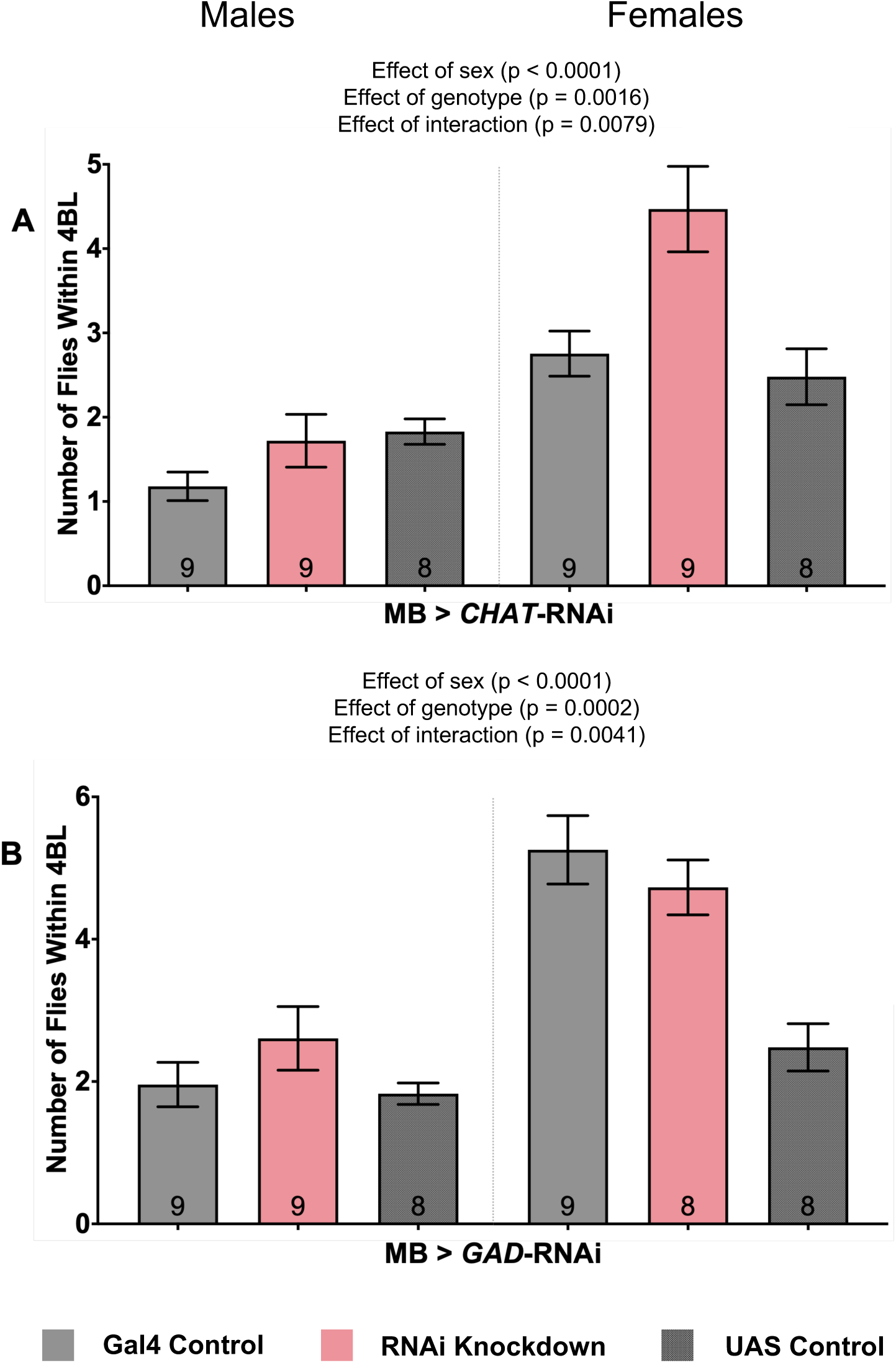
Knocking down the acetylcholine rate-limiting biosynthesis enzyme C*hAT* in the MB decreases female fly social space when looking at both genetic controls with different genetic backgrounds. Average number of flies within 4 body lengths (4BL) in male and female flies with RNAi knockdown of *ChAT* for acetylcholine synthesis (A) or *GAD1* for GABA synthesis (B) specifically in the mushroom bodies. **A.** The interaction between sex and *ChAT* knockdown shows that female flies have an increased number of flies within 4BL when compared to both the genetic control flies (two-way ANOVA – Effect of interaction: *p* = 0.0079). **B.** *GAD1* knockdown shows no difference in male and female fly social space when compared to both genetic control flies. Comparing the female *ChAT* knockdown line to both control lines shows an intermediate phenotype. Grey colour bars: genetic control *driver/+* and *Gal4/+*; pink colour bars: *driver/RNAi*. N = 8 – 9 for all treatments. Error bars are +/- s.e.m.

**Supplemental Figure 10:**
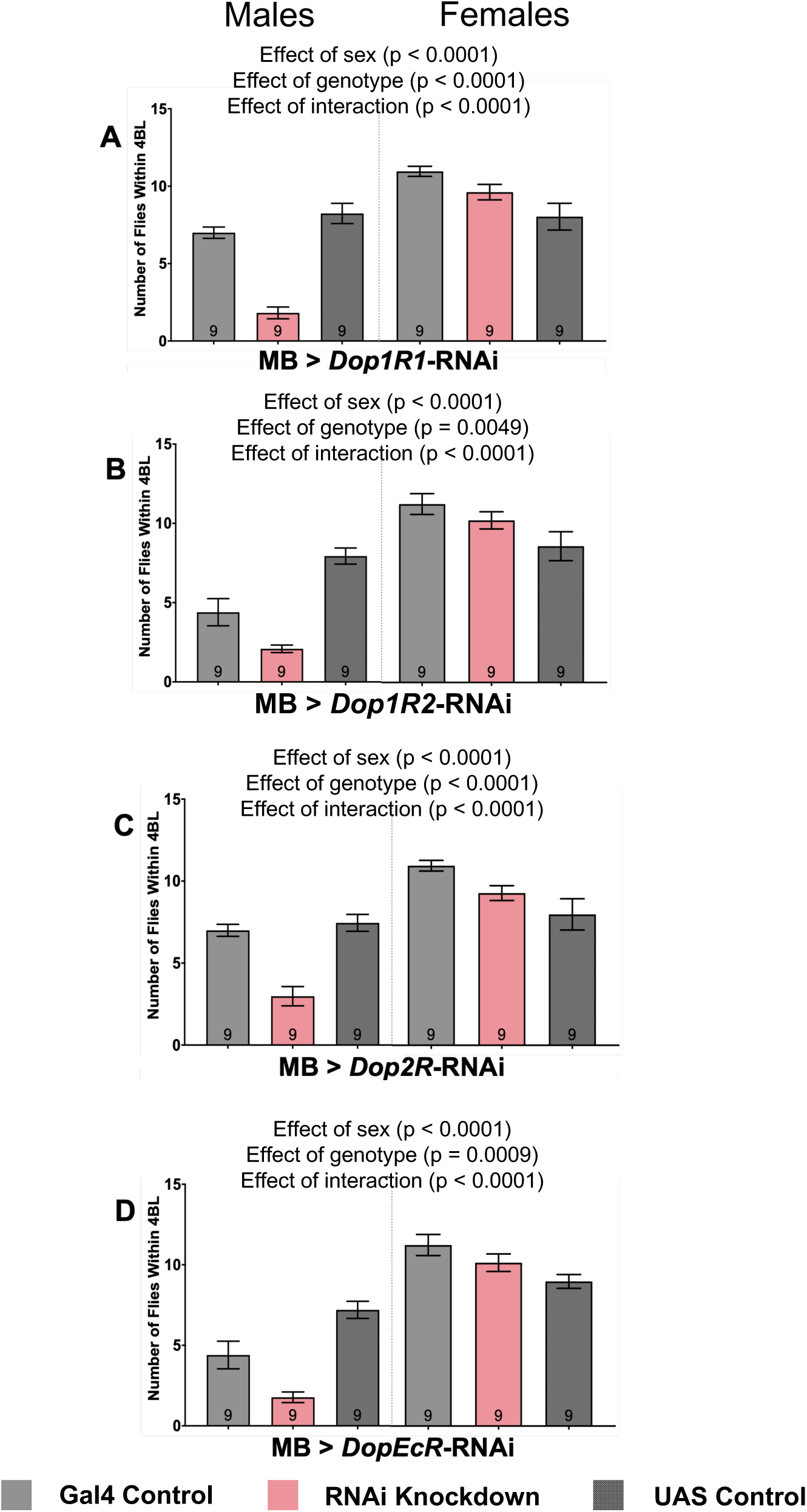
RNAi knockdown of all dopamine receptors in the MB increases male social space when looking at both genetic controls with different genetic backgrounds. The main conclusions remain unchanged when looking at both genetic controls from the V10 background even with different genetic backgrounds. Average number of flies within 4 body lengths (4BL) in male and female flies knocking down dopamine receptors *Dop1R1* (A), *Dop1R2* (B), *Dop2R* (C), and *DopEcR* (D) specifically in the mushroom bodies. **A.** *Dop1R1* knockdown in the MB of males has a decreased number of flies within 4BL compared to genetic controls (Two-way ANOVA – Effect of genotype: *F_1,32_* = 19.61, *p* < 0.0001). **B.** *Dop1R2* knockdown in the MB of males has a decreased number of flies within 4BL compared to genetic controls (Two-way ANOVA – Effect of genotype: *F_1,32_* = 41.10, *p* < 0.0001). **C.** *Dop2R* knockdown in the MB of males has a decreased number of flies within 4BL compared to genetic controls (Two-way ANOVA – Effect of genotype: *F_1,32_* = 41.10, *p* < 0.0001). **D.** *DopEcR* knockdown in the MB of males has a decreased number of flies within 4BL compared to genetic controls (Two-way ANOVA – Effect of genotype: *F_1,32_* = 8.905, *p* = 0.0054). Grey colour bars: genetic control *driver/+* and *Gal4/+*; pink colour bars: *driver/RNAi*. N = 9 for all treatments. Error bars are +/- s.e.m.

**Supplemental Figure 11:**
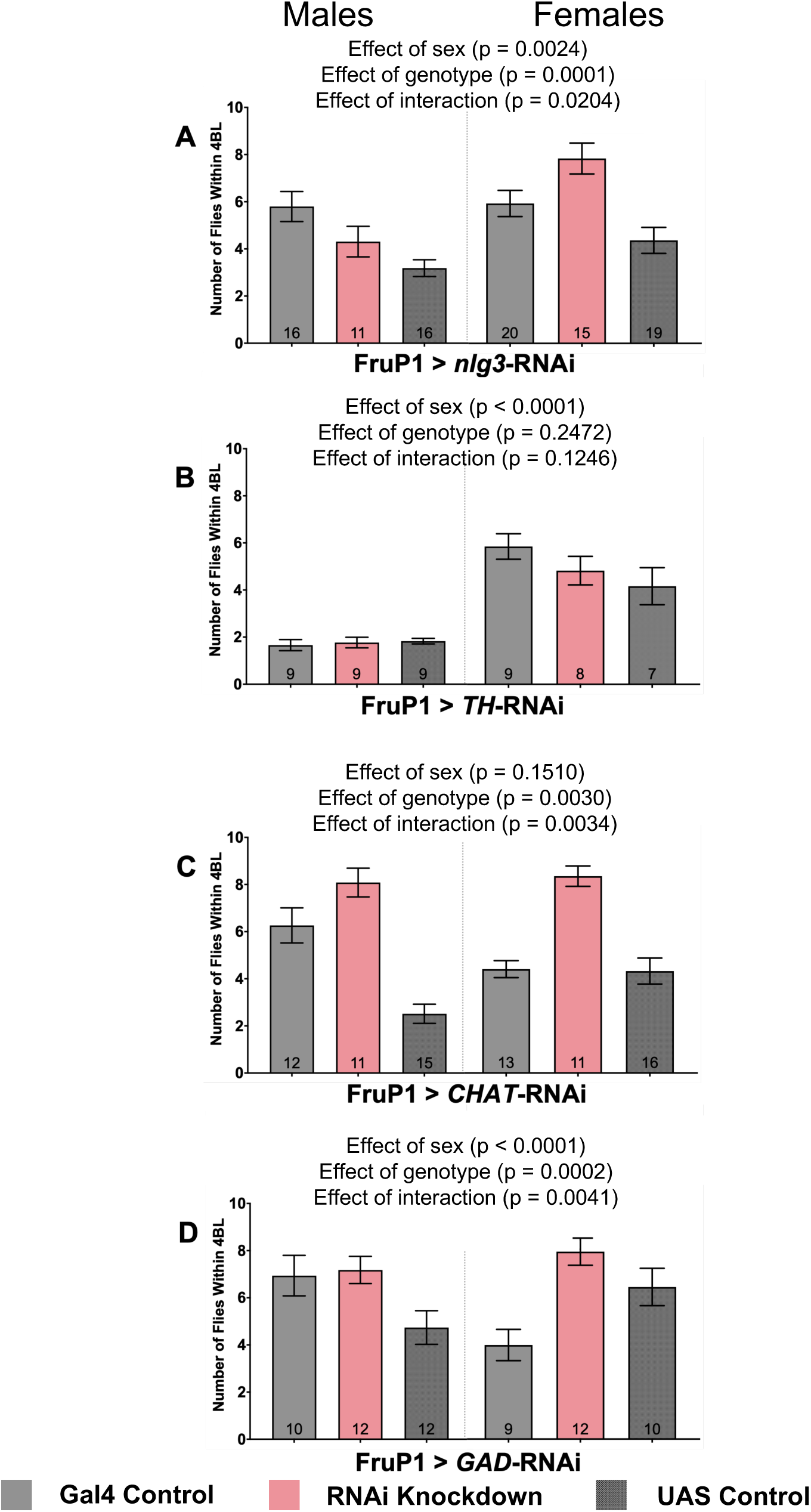
Knocking down *nlg3* in *FruP1* neurons decreases female social space, while a reduction of acetylcholine in *FruP1* neurons decreases social space in male and female flies when looking at both genetic controls with different genetic backgrounds. Average number of flies within 4 body lengths (4BL) in male and female flies with RNAi knockdown of *nlg3* (A), *TH* for dopamine synthesis (B), *ChAT* for acetylcholine synthesis (C) or *GAD1* for GABA synthesis (D) specifically in *FruP1* neurons. **A.** The interaction between sex and *nlg3* knockdown shows that female flies have a decreased number of flies within 4BL when compared to both the genetic control flies (two-way ANOVA – Effect of interaction: *p* = 0.0204). Comparing the male *nlg3* knockdown to both control lines shows an intermediate phenotype. **B.** *TH* knockdown in *FruP1* neurons does not change social space in males and females. **C.** *ChAT* knockdown in *FruP1* neurons increases the number of flies within 4BL in both males and females compared to the genetic control flies (two-way ANOVA – Effect of genotype: *p* = 0.0030). **D.** The interaction between sex and *GAD1* knockdown shows that female flies have a increased number of flies within 4BL when compared to both the genetic control flies (two-way ANOVA – Effect of interaction: *p* = 0.0041). Male *GAD1* knockdown in *FruP1* neurons shows an intermediate phenotype when compared to both genetic controls. Grey colour bars: genetic control *driver/+* and *Gal4/+*; pink colour bars: *driver/RNAi*. N = 9 - 13 for all treatments. Error bars are +/- s.e.m.

---

**Supplemental Table 1.** Genotypes of fly lines used. Fly line names, full genotypes, and sources are listed.

| Fly Line Name | Stock # | Source | Genotype | Description |
| --- | --- | --- | --- | --- |
| Control Strain |  |  |  |  |
| Cs | 64346 | Seymour Benzer | +/+ | Genetic control |
| *nlg3* Mutant |  |  |  |  |
| *MiMIC-nlg3* | 58509 | BDSC | *y[1]w[\*];Mi{y[+mDint2]=MIC}Nlg3[MI08924]* | P-element insertion in 1<sup>st</sup> intron |
| Gal4 Drivers (Outcrossed 5X in Cs) |  |  |  |  |
| *MB-Gal4* | 49265 | BDSC | *w[1118]; P{y[+t7.7] w[+mC]=GMR15E01-GAL4}attP2* | Mushroom body expression under the enhancer of *rut* |
| *PB-Gal4* | 50422 | BDSC | *w[1118]; P{y[+t7.7] w[+mC]=GMR55G08-GAL4}attP2* | Protocerebral bridge expression under the enhancer of *cmpy* |
| *OL-Gal4* | 49234 | BDSC | *w[1118]; P{y[+t7.7] w[+mC]=GMR10B01-GAL4}attP2* | Optic lobe expression under the enhancer of *crp* |
| *nlg3-Gal4* | 76134 | BDSC | *y[1] w[\*]; Mi{Trojan-GAL4.2}Nlg3[MI00445-TG4.2]* | Gal4 in *nlg3*-expressing cells |
| *fruP1-Gal4* | 66696 | BDSC | *TI{GAL4}fru[GAL4.P1.D]/TM3, Sb[1]* | Gal4 in *fruP1*-expressing cells |
| UAS Effectors |  |  |  |  |
| *TrpA1* | 26263 | BDSC | *w[\*]; P{y[+t7.7] w[+mC]=UAS-TrpA1(B).K}attP16* | Temperature control of neuron activation |
| *shit<sup>ts</sup>* | 44222 | BDSC | *w[\*]; P{w[+mC]=UAS-shit[ts1].K}3* | Temperature control of neuron silencing |
| *ChAT25-RNAi* | 25856 | BDSC | *y[1] v[1]; P{y[+t7.7] v[+t1.8]=TRiP.JF01877}attP2* | RNAi knockdown of acetylcholine receptors |
| *Gad51-RNAi* | 51794 | BDSC | *y[1] v[1]; P{y[+t7.7] v[+t1.8]=TRiP.HMC03350}attP40* | RNAi knockdown of glutamate receptors |
| *TH-RNAi* | n/a | Mark Wu (Johns Hopkins) | *w;;UAS-TH RNAi 3#G / TM6, Sb* | RNAi knockdown of dopamine receptors |

**Supplemental Table 2.** ANOVA table for all Two-Way ANOVAs performed for social space experiments presented in. **Figures 2-6**. All p values with a result less than 0.05 are bolded.

| Experiment | Figure | Effect | Df | F | p | Šidák's post hoc |  |  |
| --- | --- | --- | --- | --- | --- | --- | --- | --- |
| Social space |  |  |  |  |  | Comparison Group | Compared with | p |
| *MB>nlg3-RNAi* | 2A | **Sex**<br>**Genotype**<br>Sex * Genotype | (1,39)<br>(1,39)<br>(1,39) | **13.97**<br>**14.71**<br>4.009 | **0.0006**<br>**0.0004**<br>0.0523 | Male<br>MB>RNAi<br>Female<br>MB>RNAi | RNAi/+ | 0.3492<br>**0.0005** |
| *PB>nlg3-RNAi* | 2B | **Sex**<br>**Genotype**<br>Sex * Genotype | (1,40)<br>(1,40)<br>(1,40) | **14.90**<br>**19.02**<br>**5.607** | **0.0004**<br>**<0.0001**<br>**0.0228** | Male<br>PB>RNAi<br>Female<br>PB>RNAi | RNAi/+ | 0.2497<br>**0.0001** |
| *OL>nlg3-RNAi* | 2C | **Sex**<br>**Genotype**<br>Sex * Genotype | (1,31)<br>(1,31)<br>(1,31) | **31.29**<br>**7.191**<br>0.9424 | **<0.0001**<br>**0.0116**<br>0.3392 | Male<br>OL>RNAi<br>Female<br>OL>RNAi | RNAi/+ | 0.4048<br>**0.0320** |
| *MB>TrpA1* | 3A | **Sex**<br>**Neural Transmission**<br>Sex * Neural Transmission | (1,47)<br>(1,47)<br>(1,47) | **5.382**<br>**7.395**<br>0.172 | **0.0247**<br>**0.0091**<br>0.6796 | Male MB>Trp<br>Activated<br>Female MB>Trp<br>Activated | Inactivated | 0.2437<br>**0.0442** |
| *MB>Shi<sup>ts</sup>* | 3B | Sex<br>**Neural Transmission**<br>Sex * Neural Transmission | (1,30)<br>(1,30)<br>(1,30) | 0.680<br>**14.40**<br>0.344 | 0.4161<br>**0.0007**<br>0.5617 | Male MB>Shi<br>Activated<br>Female MB>Shi<br>Activated | Inactivated | **0.0105**<br>0.0518 |
| *PB>TrpA1* | 3C | Sex<br>Neural Transmission<br>Sex * Neural Transmission | (1,74)<br>(1,74)<br>(1,74) | 39.94<br>17.72<br>13.62 | 0.0868<br>0.2018<br>0.1953 | Male PB>Trp<br>Activated<br>Female PB>Trp<br>Activated | Inactivated | 0.9999<br>0.1382 |
| *PB>Shi<sup>ts</sup>* | 3D | Sex<br>Neural Transmission<br>Sex * Neural Transmission | (1,31)<br>(1,31)<br>(1,31) | 0.6335<br>3.443<br>1.483 | 0.4321<br>0.0730<br>0.2325 | Male PB>Shi<br>Activated<br>Female PB>Shi<br>Activated | Inactivated | 0.8897<br>0.0596 |
| *Trojan>TrpA1* | 3E | Sex<br>**Neural Transmission**<br>Sex * Neural Transmission | (1,28)<br>(1,28)<br>(1,28) | 2.380<br>**12.70**<br>1.366 | 0.1341<br>**0.0013**<br>0.2523 | Male Trojan>Trp<br>Activated<br>Female Trojan>Trp<br>Activated | Inactivated | 0.2214<br>**0.0029** |
| *Trojan>Shi<sup>ts</sup>* | 3F | **Sex**<br>**Neural Transmission**<br>Sex * Neural Transmission | (1,22)<br>(1,22)<br>(1,22) | **4.666**<br>**5.998**<br>**6.169** | **0.0419**<br>**0.0228**<br>**0.0211** | Male MB>Trp<br>Activated<br>Female MB>Trp<br>Activated | Inactivated | 0.9996<br>**0.0042** |
| *MB>CHAT25-RNAi* | 4A | **Sex**<br>**Genotype**<br>Sex * Genotype | (1,32)<br>(1,32)<br>(1,32) | **41.09**<br>**11.19**<br>3.020 | **<0.0001**<br>**0.0021**<br>0.0919 | Male<br>MB>RNAi<br>Female<br>MB>RNAi | RNAi/+ | 0.4588<br>**0.0022** |
| *MB>GAD51-RNAi* | 4B | **Sex**<br>**Genotype**<br>Sex * Genotype | (1,29)<br>(1,29)<br>(1,29) | **15.22**<br>**18.10**<br>**4.276** | **0.0005**<br>**0.0002**<br>**0.0477** | Male<br>MB>RNAi<br>Female<br>MB>RNAi | RNAi/+ | 0.2389<br>**0.0003** |
| *MB>Dop1R1-RNAi* | 5A | **Sex**<br>**Genotype**<br>Sex * Genotype | (1,32)<br>(1,32)<br>(1,32) | **65.51**<br>**19.61**<br>**27.24** | **<0.0001**<br>**<0.0001**<br>**<0.0001** | Male<br>MB>RNAi<br>Female<br>MB>RNAi | MB/+ | **<0.0001**<br>0.0505 |
| *MB>Dop1R2-RNAi* | 5B | **Sex**<br>**Genotype**<br>Sex * Genotype | (1,32)<br>(1,32)<br>(1,32) | **139.4**<br>**7.401**<br>1.168 | **<0.0001**<br>**0.0105**<br>0.2880 | Male<br>MB>RNAi<br>Female<br>MB>RNAi | MB/+ | **0.0225**<br>0.4447 |
| *MB>Dop2R-RNAi* | 5C | **Sex**<br>**Genotype**<br>Sex * Genotype | (1,32)<br>(1,32)<br>(1,32) | **120.7**<br>**41.10**<br>**7.448** | **<0.0001**<br>**<0.0001**<br>**0.0102** | Male<br>MB>RNAi<br>Female<br>MB>RNAi | MB/+ | **<0.0001**<br>**0.0276** |
| *MB>DopEcR-RNAi* | 5D | **Sex**<br>**Genotype**<br>Sex * Genotype | (1,32)<br>(1,32)<br>(1,32) | **139.3**<br>**8.905**<br>1.586 | **<0.0001**<br>**0.0054**<br>0.2167 | Male<br>MB>RNAi<br>Female<br>MB>RNAi | MB/+ | **0.0103**<br>0.4098 |
| *FruP1>TrpA1* | 6A | **Sex**<br>**Neural Transmission**<br>Sex * Neural Transmission | (1,33)<br>(1,33)<br>(1,33) | **5.168**<br>**22.65**<br>0.6987 | **0.0296**<br>**<0.0001**<br>0.4092 | Male FruP1>Trp<br>Activated<br>Female FruP1>Trp<br>Activated | Inactivated | **0.0241**<br>**0.0004** |
| *FruP1>Shi<sup>ts</sup>* | 6B | Sex<br>Neural Transmission<br>Sex * Neural Transmission | (1,30)<br>(1,30)<br>(1,30) | 0.5671<br>2.350<br>3.451 | 0.4573<br>0.1358<br>0.0730 | Male FruP1>Trp<br>Activated<br>Female FruP1>Trp<br>Activated | Inactivated | **0.0453**<br>0.9676 |
| *FruP1>nlg3-RNAi* | 7A | **Sex**<br>**Genotype**<br>Sex * Genotype | (1,57)<br>(1,57)<br>(1,57) | **15.57**<br>**16.27**<br>**4.090** | **0.0002**<br>**0.0002**<br>**0.0478** | Male<br>FruP1>RNAi<br>Female<br>FruP1>RNAi | RNAi/+ | 0.3363<br>**<0.0001** |
| *FruP1>TH-RNAi* | 7B | **Sex**<br>**Genotype**<br>Sex * Genotype | (1,29)<br>(1,29)<br>(1,29) | **32.36**<br>0.413<br>0.603 | **<0.0001**<br>0.5255<br>0.4436 | Male<br>FruP1>RNAi<br>Female<br>FruP1>RNAi | RNAi/+ | 0.9938<br>0.5709 |
| *FruP1>CHAT25-RNAi* | 7C | Sex<br>**Genotype**<br>Sex * Genotype | (1,47)<br>(1,47)<br>(1,47) | 16.12<br>**99.11**<br>**0.7134** | 0.0002<br>**<0.0001**<br>**0.4026** | Male<br>FruP1>RNAi<br>Female<br>FruP1>RNAi | RNAi/+ | **<0.0001**<br>**<0.0001** |
| *FruP1>GAD51-RNAi* | 7D | **Sex**<br>**Genotype**<br>Sex * Genotype | (1,42)<br>(1,42)<br>(1,42) | **11.81**<br>**8.478**<br>0.2264 | **0.0013**<br>**0.0057**<br>0.6367 | Male<br>FruP1>RNAi<br>Female<br>FruP1>RNAi | RNAi/+ | **0.0363**<br>0.1897 |

**Supplemental Table 3.**
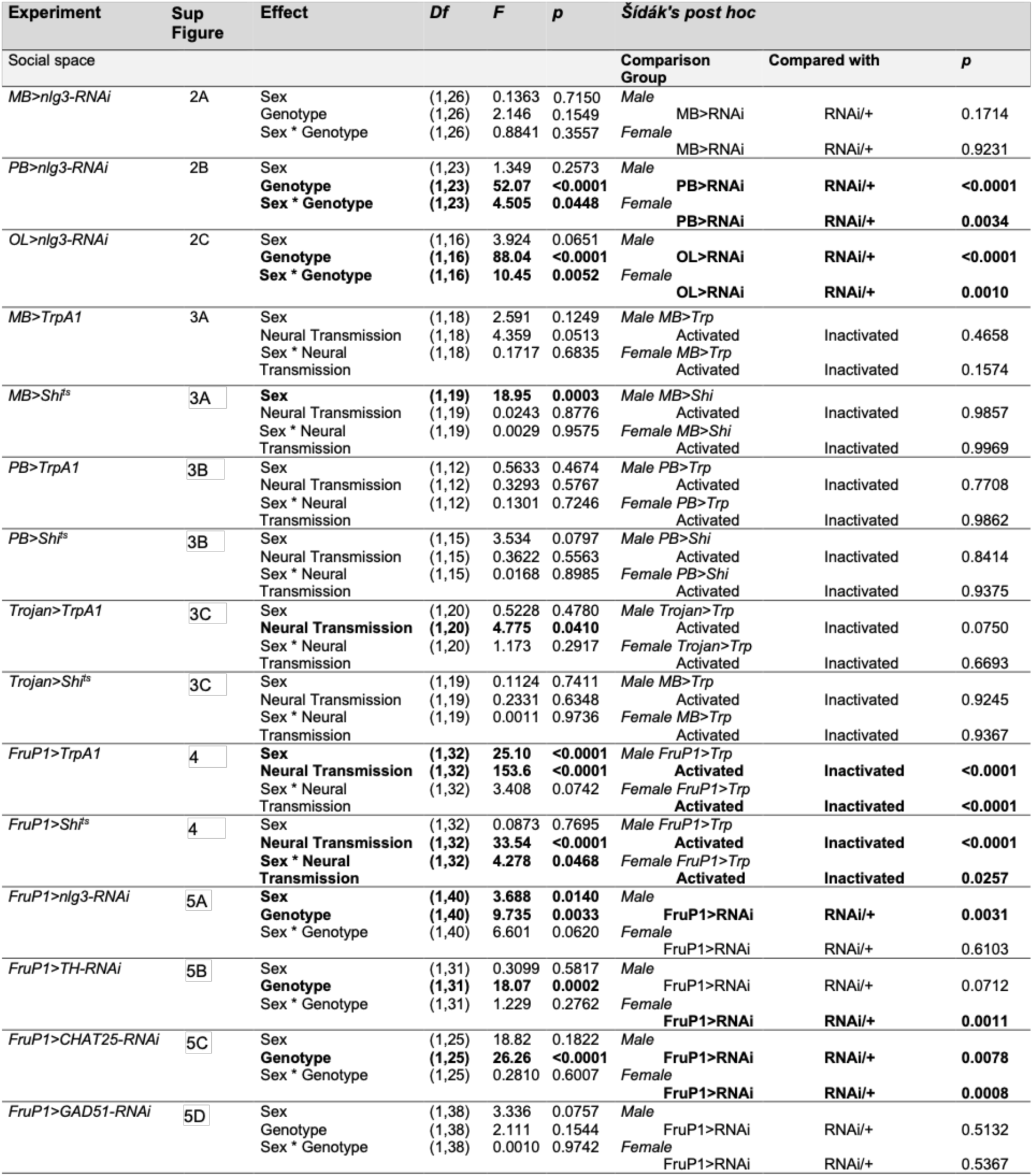
ANOVA table for all Two-Way ANOVAs performed for climbing experiments presented in Supplemental Figures 2-5. All p values with a result less than 0.05 are bolded.

